# Cardelino: Integrating whole exomes and single-cell transcriptomes to reveal phenotypic impact of somatic variants

**DOI:** 10.1101/413047

**Authors:** Davis J. McCarthy, Raghd Rostom, Yuanhua Huang, Daniel J. Kunz, Petr Danecek, Marc Jan Bonder, Tzachi Hagai, HipSci Consortium, Wenyi Wang, Daniel J. Gaffney, Benjamin D. Simons, Oliver Stegle, Sarah A. Teichmann

## Abstract

Decoding the clonal substructures of somatic tissues sheds light on cell growth, development and differentiation in health, ageing and disease. DNA-sequencing, either using bulk or using single-cell assays, has enabled the reconstruction of clonal trees from frequency and co-occurrence patterns of somatic variants. However, approaches to systematically characterize phenotypic and functional variations between individual clones are not established. Here we present cardelino (https://github.com/PMBio/cardelino), a computational method for inferring the clone of origin of individual cells that have been assayed using single-cell RNA-seq (scRNA-seq). After validating our model using simulations, we apply cardelino to matched scRNA-seq and exome sequencing data from 32 human dermal fibroblast lines, identifying hundreds of differentially expressed genes between cells from different somatic clones. These genes are frequently enriched for cell cycle and proliferation pathways, indicating a key role for cell division genes in non-neutral somatic evolution.

**Key findings:**

- A novel approach for integrating DNA-seq and single-cell RNA-seq data to reconstruct clonal substructure for single-cell transcriptomes.
- Evidence for non-neutral evolution of clonal populations in human fibroblasts.
- Proliferation and cell cycle pathways are commonly distorted in mutated clonal populations.

## Introduction

Ageing, environment and genetic factors can impact mutational processes, thereby shaping the acquisition of somatic mutations across the life span (Burnet, 1974; Martincorena and Campbell, 2015; Stransky *et al*., 2011; Hodis *et al*., 2012; Huang *et al*., 2018). The maintenance and evolution of somatic mutations in different sub-populations of cells can result in clonal structure, both within healthy and disease tissues. Targeted, whole-genome and whole-exome DNA sequencing of bulk cell populations has been utilized to reconstruct the mutational processes that underlie somatic mutagenesis (Nik-Zainal *et al*., 2012; Alexandrov *et al*., 2013; Forbes *et al*., 2017; Bailey *et al*., 2018; Ding *et al*., 2018) as well as clonal trees (Roth *et al*., 2014; Deshwar *et al*., 2015; Jiang *et al*., 2016).

Availability of single-cell DNA sequencing methods (scDNA-seq; Navin *et al*., 2011; Wang *et al*., 2014; Navin, 2015) combined with new computational approaches have helped to improve the reconstruction of clonal populations (Kim and Simon, 2014; Navin and Chen, 2016; Jahn *et al*., 2016; Kuipers *et al*., 2017; Roth *et al*., 2016; Salehi *et al*., 2017; Malikic *et al*., 2017). However, the functional differences between clones and their molecular phenotypes remain largely unknown. Systematic characterisation of the phenotypic properties of clones could reveal mechanisms underpinning healthy tissue growth and the transition from normal to malignant behaviour.

An important step towards such functional insights would be access to genome-wide expression profiles of individual clones, yielding genotype-phenotype connections for clonal architectures in tissues. Recent studies have explored mapping scRNA-seq profiles to clones with distinct copy number states in cancer, thus providing a first glimpse at clone-to-clone gene expression differences in disease (Müller *et al*., 2016; Tirosh *et al*., 2016; Fan *et al*., 2018). However, generally-applicable methods for inferring the clone of origin of single cells to study genotype-transcriptome relationships are not yet established.

To address this, we have developed cardelino: a computational method that exploits variant information in scRNA-seq reads to map cells to their clone of origin. After validating our model using simulations, we demonstrate that cardelino allows for accurate assignment of full-length single-cell transcriptomes to the clonal substructure in 32 normal dermal fibroblast lines. With linked somatic variants, clone and gene expression information, we investigate gene expression differences between clones at the level of individual genes and in pathways, which provides new insights into the dynamics of clones. These findings also extend recent studies using bulk DNA-seq data, predominantly in epithelial cells, that have revealed oncogenic mutations and evidence of selective clonal dynamics in normal tissue samples (*Behjati *et al*., 2014;* Martincorena *et al*., 2015; Simons, 2016b; Martincorena et al., 2016; Simons, 2016a). Our approach can be applied to a broad range of somatic substructure analyses in population or disease settings to reveal previously inaccessible differences in molecular phenotypes between cells from the same individual.

## Results

### Mapping single-cell transcriptomes to somatic clones with cardelino

We present a two-step strategy for integrating somatic clonal substructure and transcriptional heterogeneity within a population of cells. First, a clonal tree is inferred using variant allele frequencies from bulk or single-cell DNA sequencing data (Roth *et al*., 2014; Deshwar *et al*., 2015; Jiang *et al*., 2016; Kim and Simon, 2014; Navin and Chen, 2016; Jahn *et al*., 2016; Kuipers *et al*., 2017; Roth *et al*., 2016; Salehi *et al*., 2017; Malikic *et al*., 2017). Subsequently, cardelino performs probabilistic assignment of individual single-cell transcriptomes to inferred clones, using variant information extracted from single-cell RNA-seq reads (**Fig. 1a**).

**Figure 1.**
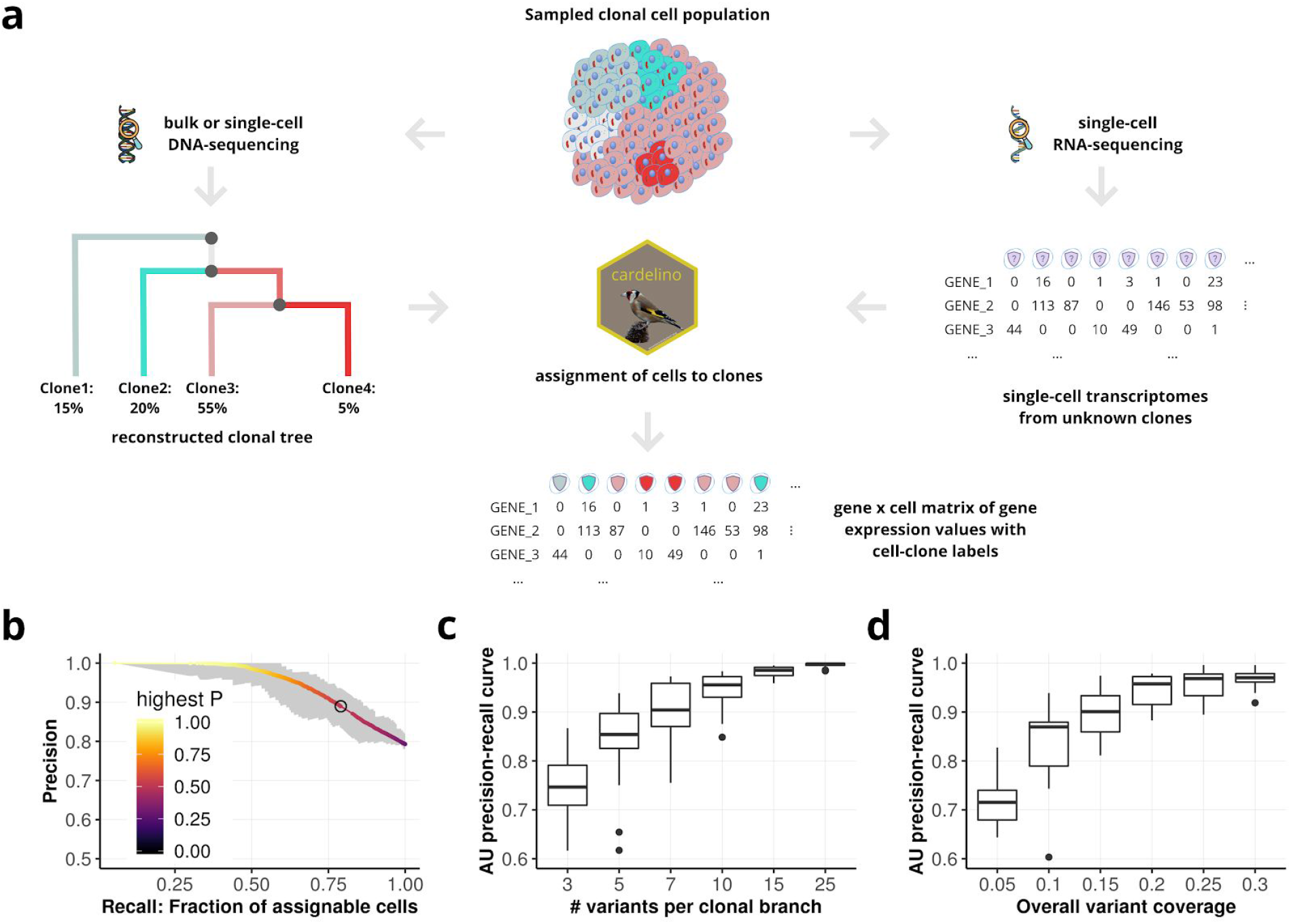
Overview and validation of the cardelino model. **(a)** Overview and approach. A clonal tree is reconstructed based on DNA-sequencing (e.g. deep exome sequencing) data, followed by probabilistic mapping of single-cell transcriptomes to reconstructed clones based on variants detected in scRNA-seq reads. **(b-d)** Benchmark of the cell assignment step using simulations. **(b)** Overall assignment performance for 200 cells, when simulating from 4-clone structures with 10 variants per branch and non-zero read coverage for 20% of variants (**Methods**). Shown is the fraction of true positive cell assignments (precision) as a function of the fraction of assigned cells (recall) when varying the threshold of the cell assignment probability. Shaded areas denote upper and lower quartiles from 20 repeat experiments. The black circle corresponds to the posterior P threshold of 0.5. **(c,d)** Area Under (AU) precision-recall curve *(i.e.* area under curves such as shown in **b**), when varying the numbers of variants per clonal branch (**c**) and the average fraction of detectable clone-specific variants (non-zero scRNA-seq read coverage) (**d**). For details and default parameter settings see **Methods**.

Briefly, cardelino is based on a Bayesian mixture model, where individual mixture components correspond to somatic clones in a given population of cells. To enable accurate cell assignment, we employ a beta-binomial error model that accounts for dropout and allelic imbalance, parameters that affect the sensitivity for detecting true somatic variants from sparse scRNA-seq data. Initially, we assess the accuracy of our approach using simulated data that mimic typical clonal structures and properties of scRNA-seq as observed in real data (4 clones, 10 variants per branch, 20% of variants with read coverage, 200 cells, 20 repeat experiments; **Methods**). Cardelino achieves high overall performance (Precision-Recall AUC=0.947; **Fig. 1b**); for example, at a threshold of P=0.5 for the cell assignment confidence (posterior probability of cell assignment), the model assigns 79% of all cells with 89.0% accuracy.

We explore the effect of key dataset characteristics on cell assignment, including the number of mutations per clonal branch (**Fig. 1c**), the expected number of variants with non-zero scRNA-seq coverage per cell (**Fig. 1d**), the number of clones and the overall extent and variability of allelic imbalance across genes (**Methods; Supp. Fig. S1**). We also assess individual modelling assumptions in cardelino, confirming the expected benefits from a full Bayesian treatment and the explicit error model for allelic dropout as used in cardelino (**Fig. S1**).

Taken together, these results indicate that cardelino is broadly applicable to robustly assign individual single-cell transcriptomes to clones, thereby reconstructing clone-specific transcriptome profiles.

### Cardelino assigns single cell transcriptomes to clones in human dermal fibroblasts

Next, we apply cardelino to 32 human dermal fibroblast lines derived from healthy donors that are part of the UK human induced pluripotent stem cell initiative (HipSci; Kilpinen *et al*., 2017). For each line, we generated deep whole exome sequencing data (WES; median read coverage: 254), and matched Smart-seq2 scRNA-seq profiles using pools of three lines in each processing batch (**Methods**). We assayed between 30 and 107 cells per line (median 61 cells after QC; median coverage: 484k reads; median genes observed: 11,108; **Supp. Table S2**).

Initially, we consider high-confidence somatic single nucleotide variants (SNVs) identified based on WES data (**Methods**) to explore the mutational landscape across lines. This reveals considerable variation in the total number of somatic SNVs, with 41–612 variants per line (**Fig. 2a**). The majority of SNVs can be attributed to the well-documented UV signature, COSMIC Signature 7 (primarily C to T mutations; Forbes *et al*., 2017), agreeing with expected mutational patterns from UV exposure of skin tissues (**Fig. 2a; Supp. Fig. S2; Methods**).

**Figure 2.**
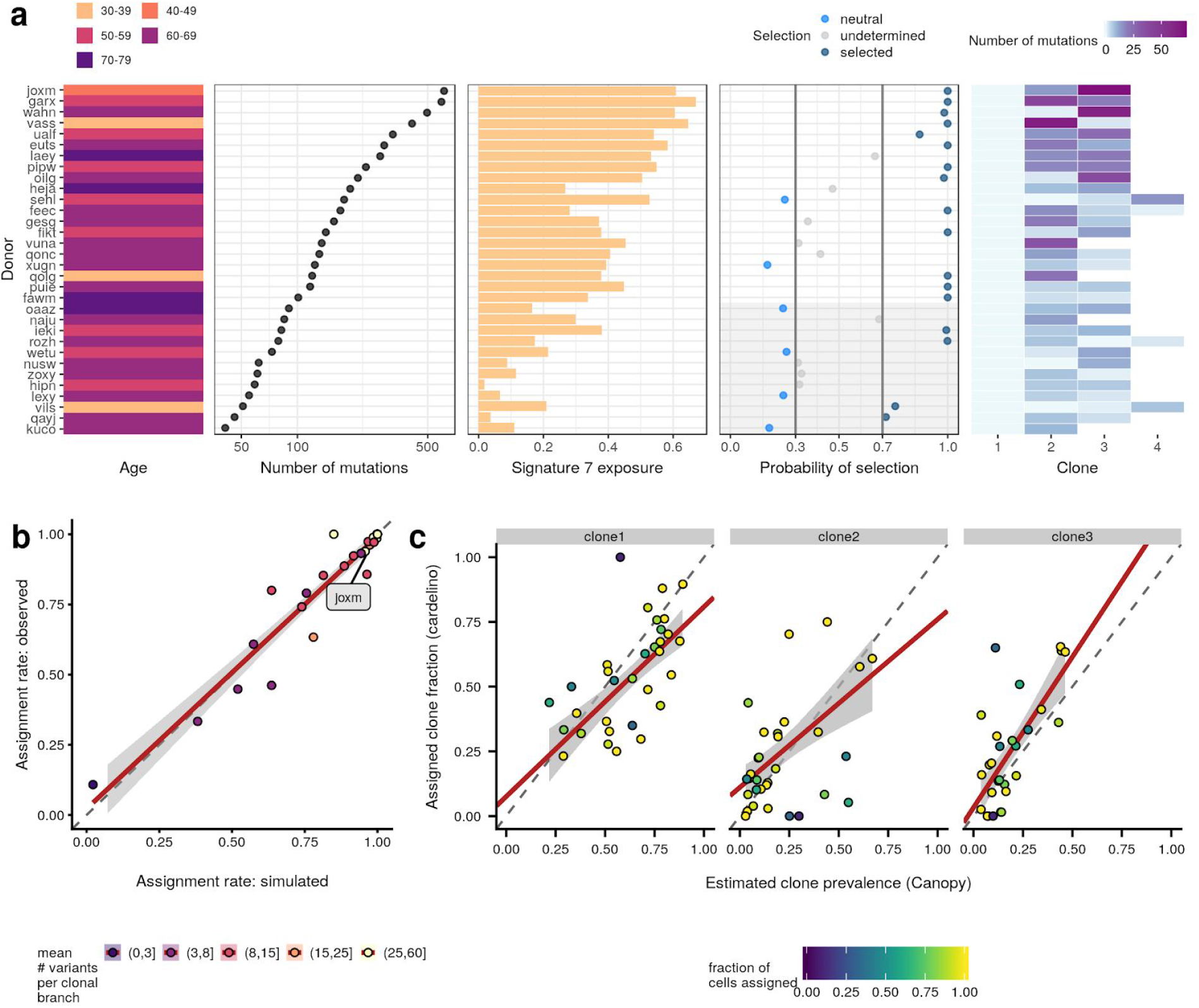
Parallel deep exome sequencing and scRNA-seq profiling of 32 human dermal fibroblast lines. (a) Overview and somatic mutation profiles across lines (donors), from left to right: donor age; number of somatic SNVs; estimated exposure of COSMIC mutational signature 7; probability of selection estimated by SubClonalSelection (Williams *et al*., 2018), colour denotes the selection status based on probability cut-offs (grey lines), the grey background indicates results with high uncertainty due to the low number of mutations detected; number of clones inferred using Canopy (Jiang et al., 2016), with colour indicating the number of informative somatic SNVs for cell assignment to each clone (non-zero read coverage in scRNA-seq data). **(b)** Assignment rate (fraction of cells assigned) using simulated single-cell transcriptomes (x-axis; **Methods**) *versus* the empirical assignment rate (y-axis) for each line (at assignment threshold posterior P>0.5). Colour denotes the average number of informative variants across clonal branches per line. The line-of-best fit from a linear model is shown in red, with 95% confidence interval shown in grey. **(c)** Estimated clone prevalence from WES data (x-axis; using Canopy) versus the fraction of single-cell transcriptomes assigned to the corresponding clone (y-axis; using cardelino). Shown are the fractions of cells assigned to clones one to three as in **a**, considering the most likely assignment for assignable cells (posterior probability P>0.5) with each point representing a cell line; see **Supp. Fig. S11** for results from four donors with >3 clones). Colour denotes the total fraction of assignable cells per line (P>0.5). A line-of-best fit from a weighted regression model is shown in red with 95% confidence interval shown in grey.

To understand whether the somatic SNVs confer any selective advantage in skin fibroblasts, we used SubConalSelection to identify neutral and selective dynamics at a per-donor level (Williams *et al*., 2018). Other established methods such as dN/dS (Martincorena *et al*., 2018) and alternative methods using the SNV frequency distribution (Simons, 2016a; Williams *et al*., 2016) are not conclusive in the context of this dataset due to lack of statistical power resulting from the low number of mutations detected in each sample. The SubClonalSelection analysis identifies at least 10 lines with a clear fit to their selection model, suggesting positive selection of clonal sub-populations (**Fig. 2a; Supp. Fig. 3; Methods**). In other words, a third of the samples from this cohort of healthy donors contain clones evolving adaptively, which we can investigate in more detail in terms of transcriptome phenotype.

Next, we reconstruct the clonal trees in each line using WES-derived estimates of the variant allele frequency of somatic variants that are also covered by scRNA-seq reads (**Methods**). Canopy (Jiang *et al*., 2016) identifies two to four clones per line (**Fig. 2a**). Following this, we use cardelino to map scRNA-seq profiles from a total of 1,728 cells to clones from the corresponding lines (**Methods**), with an assignment rate of 33–100% (at posterior probability P>0.5; corresponding to 4 to 107 cells; median 49.5 cells).

To assess the confidence of these cell assignments, we consider simulated cells drawn from a clonal structure that matches the corresponding line, observing high concordance (*R* ^2^ = 0.92) between the expected and empirical cell-assignment rates (**Fig. 2b**). Lines with clones that harbour fewer distinguishing variants are associated with lower assignment rates (**Supp. Fig. S4**), at consistently high cell assignment accuracy (median 0.976, mean 0.950; **Supp. Fig. S5**), indicating that the posterior probability of assignment is calibrated across different settings. We also consider the impact of technical features of scRNA-seq data on cell assignment, finding no evidence of biased cell assignments (**Supp. Fig. 6-10**). Finally, clone prevalences estimated from Canopy and the fractions of cells assigned to the corresponding clones are markedly concordant (overall *R* ^2^ = 0.61), providing additional confidence in the cardelino cell assignments (**Fig. 2c**).

### Differences in gene expression between clones suggest phenotypic impact of somatic mutations

Initially, we focus on the fibroblast line with the largest number of somatic SNVs (*joxm;* white female aged 45-49; **Fig. 2a**), with 612 somatic SNVs (112 detected both in WES and scRNA-seq) and 79 QC-passing cells, 97% of which could be assigned to one of three clones (**Fig. 3a**). Principal component analysis of the scRNA-seq profiles of these cells reveals global transcriptome substructure that is aligned with the somatic clonal structure in this population of cells (**Fig. 3b**). Additionally, we observe differences in the fraction of cells in different cell cycle stages, where clone1 has the fewest cells in G1, and the largest fraction in S and G2M (**Fig. 3b, inset plot;** PC1 in **Supp. Fig. 12-13**). This suggests that clone1 is proliferating most rapidly. Next, we consider differential expression analysis of individual genes between the two largest clones (clone1: 45 cells *versus* clone2: 25 cells), which identifies 446 DE genes (edgeR QL F-test; FDR<0.1; **Fig. 3c**). These genes are approximately evenly split into up/down-regulated sets. However, the downregulated genes are enriched for processes involved in the cell cycle and cell proliferation. Specifically, the three significantly enriched gene sets are all up-regulated in clone1 (camera; FDR<0.05; **Fig. 3d**). All three gene sets (E2F targets, G2M checkpoint and mitiotic spindle) are associated with the cell cycle, so these results are consistent with the cell cycle stage assignments suggesting increased proliferation of clone1.

**Figure 3.**
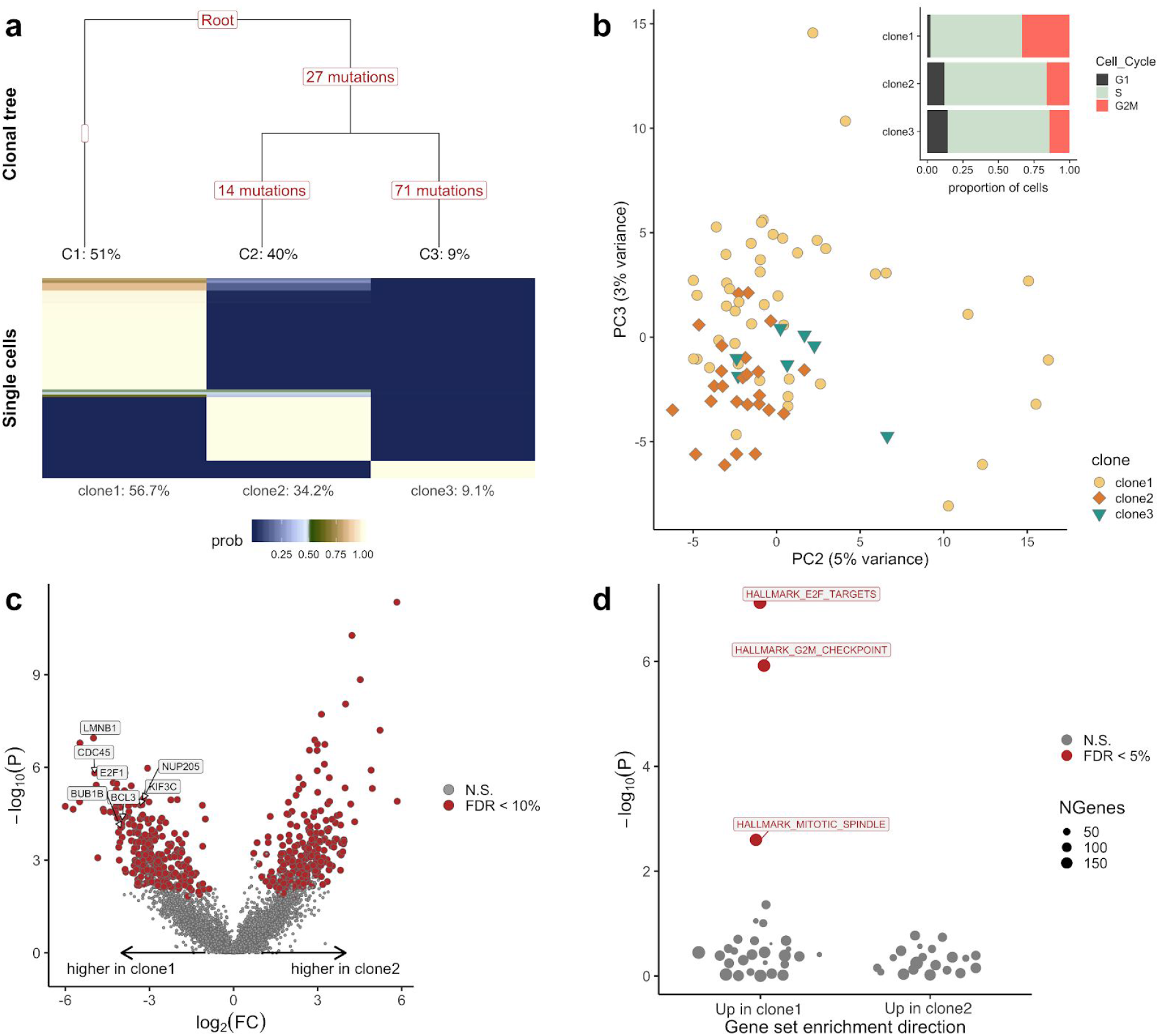
Clone-specific transcriptome profiles reveal gene expression differences for *joxm*, one example line. **(a)** Top: Clonal tree inferred using Canopy (Jiang et al., 2016). The number of variants tagging each branch and the expected prevalence (fraction) of each clone is shown. Bottom: cardelino cell assignment matrix, showing the assignment probability of individual cells to three clones. Shown below each clone is the fraction of cells assigned to each clone. **(b)** Principal component analysis of scRNA-seq profiles with colour indicating the most likely clone assignment. Inset plot: Cell-cycle phase fractions for cells assigned to each clone (using cyclone; Scialdone et al., 2015). **(c)** Volcano plot showing negative log_10_ P values versus log fold changes (FC) for differential expression between cells assigned to clone2 and clone1. Significant differentially expressed genes (FDR<0.1) are highlighted in red. **(d)** Enrichment of MSigDB Hallmark gene sets using camera (Wu and Smyth, 2012) based on log_2_ FC values between clone2 and clone1 as in **c**. Shown are negative log_10_ P values of gene set enrichments, considering whether gene sets are up-regulated in clone1 or clone2, with significant (FDR < 0.05) gene sets highlighted and labelled. All results are based on 77 out of 79 cells that could be confidently assigned to one clone (posterior P>0.5; **Methods**).

Taken together, the results suggest that somatic substructure in this cell population results in clones that exhibit measurably different expression phenotypes across the transcriptome, with significant differential expression in cell cycle and growth pathways.

### Cell cycle and proliferation pathways frequently vary between clones

To quantify the overall effect of somatic substructure on gene expression variation across the entire dataset, we fit a linear mixed model to individual genes (**Methods**), partitioning gene expression variation into a line (likely donor) component, a clone component, technical batch (*i.e*. processing plate), cellular detection rate (proportion of genes with non-zero expression per cell) and residual noise. As expected, the line component typically explains a substantially larger fraction of the expression variance than clone (median 5.5% for line, 0.5% for clone), but there are 194 genes with a substantial clone component (>5% variance explained by clone; **Fig. 4a**). Even larger clone effects are observed when estimating the clone component in each line separately, which identifies between 331 and 2,162 genes with a substantial clone component (>5% variance explained by clone; median 825 genes; **Fig. 4b**). This indicates that there are line-specific differences in the set of genes that vary with clonal structure.

**Figure 4.**
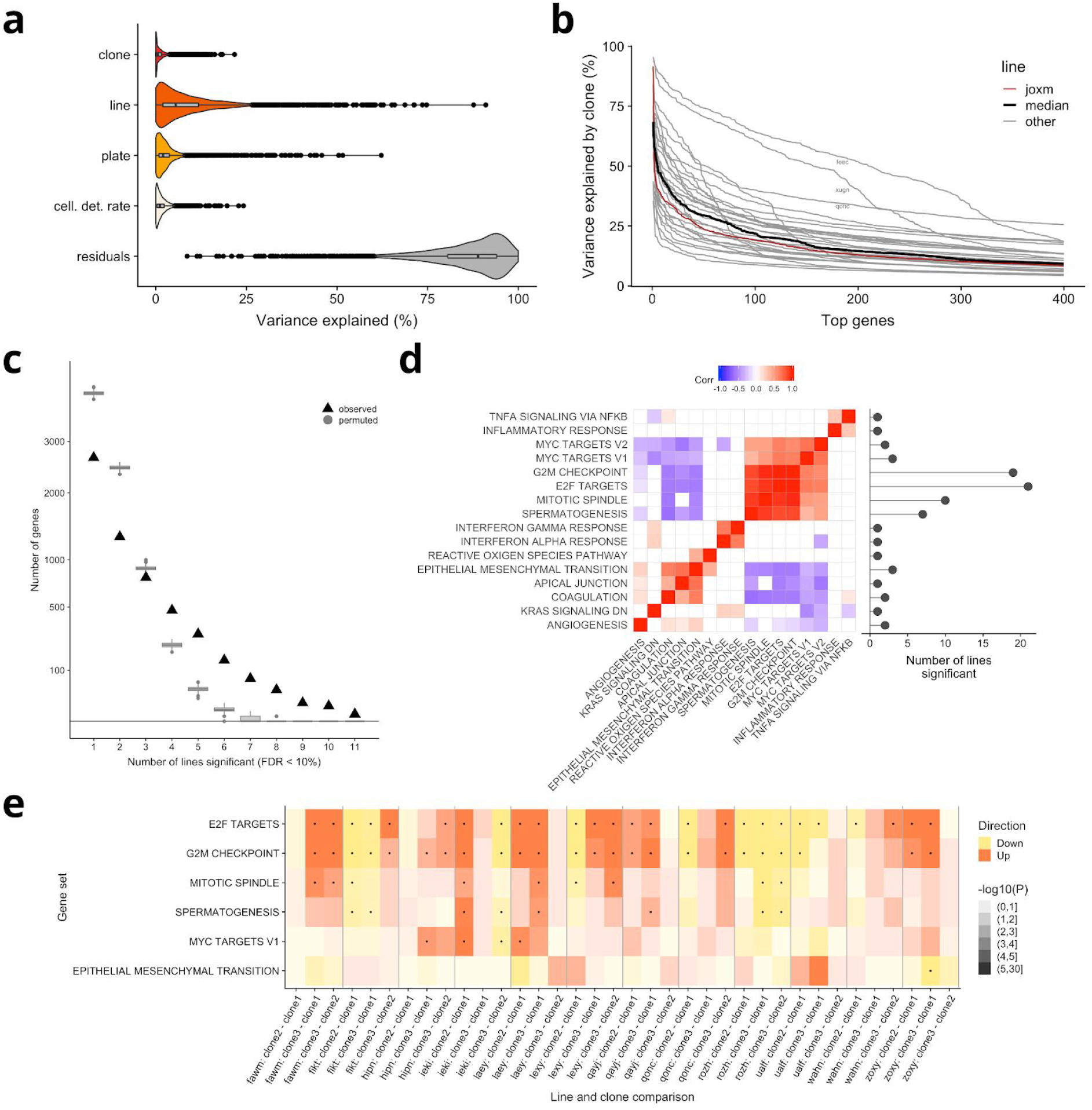
Signatures of transcriptomic clone-to-clone variation across 31 lines. **(a)** Violin and box plots show the percentage of variance explained by clone, line, experimental plate and cellular detection rate for 4,998 highly variable genes, estimated using a linear mixed model (**Methods**). **(b)** Percentage of gene expression variance explained by clone when fitting a linear mixed model for each individual line for the 400 genes with the most variance explained by clone per line (**Methods**). Individual lines correspond to cell lines (donors), with *joxm* highlighted in red and the median across all lines in black. **(c)** The number of recurrently differentially expressed (DE) genes between any pair of clones (FDR<0.1; edgeR QL F test), detected in at least one to 11 lines, with box plots showing results expected by chance (using 1,000 permutations). **(d)** Left panel: Heatmap showing pairwise correlation coefficients (Spearman R, only nominal significant correlations shown (P<0.05)) between signed P-values of gene set enrichment across donors, based on differentially expressed genes between clones. Shown are the 16 most frequently enriched MSigDB Hallmark gene sets. Right panel: number of lines in which each gene set is found to be significantly enriched (FDR<0.05). **(e)** Heatmap depicting signed P-values of gene set enrichments for six Hallmark gene sets in 12 lines. Dots denote significant enrichments (FDR<0.05).

Next, we carry out a systematic differential expression (DE) analysis to assess transcriptomics differences between any pair of clones for each line (considering 31 lines with at least 15 cells for DE testing; **Methods**). This identifies up to 2,401 DE genes per line (FDR<0.1, edgeR QL F test). A majority, 52%, of the total set of 5,798 unique DE genes, are detected in two or more lines, and 31% are detected in at least three of the 31 lines. Comparison to data with permuted gene labels demonstrates an excess of recurrently differentially expressed genes compared to chance expectation (**Fig 4c**, P<0.001; 1,000 permutations; **Methods**). We also identify a small number of genes that contain somatic variants in a subset of clones, resulting in differential expression between wild-type and mutated clones (**Supp. Fig. S14**).

To investigate the transcriptomic changes between cells in more detail, we use gene set enrichment analysis of DE genes in each line. This reveals whether there is functional convergence at a pathway level (using MSigDB Hallmark gene sets; **Methods**; Liberzon *et al*., 2011). Of 31 lines tested, 21 have at least one significant MSigDB Hallmark gene set (FDR<0.05, camera; **Methods**), with key gene sets related to cell cycle and growth being significantly enriched in all of those 21 lines. Directional gene expression changes of gene sets for the *E2F* targets, G2M checkpoint, mitotic spindle and MYC target pathways are highly coordinated (**Fig. 4d**), despite limited overlap of individual genes between the gene sets (**Supp. Fig. S15**).

Similarly, directional expression changes for pathways of epithelial-mesenchymal transition (EMT) and apical junction are correlated with each other. Interestingly, these are anti-correlated with expression changes in cell cycle and proliferation pathways (**Fig. 4d**). Within individual lines, the enrichment of pathways often differs between pairs of clones, highlighting the variability in effects of somatic variants on the phenotypic behaviour of cells (**Fig. 4e;** all lines shown in **Supp. Fig. S16**).

These consistent pathway enrichments across a larger set of donors point to somatic mutations commonly affecting the cell cycle and cell growth in fibroblast cell populations. These results indicate both deleterious and adaptive effects of somatic variants on proliferation, suggesting that a significant fraction of these variants are non-neutral in the majority of donors in our study.

## Discussion

Here, we develop and apply a computational approach for integrating somatic clonal structure with single-cell RNA-seq data This allows us to identify molecular signatures that differ between clonal cell populations. Our approach is based on first inferring clonal structure in a population of cells using WES data, followed by the assignment of individual single-cell transcriptomes using a computational approach called cardelino. Our method enables the efficient reconstruction of clone-specific transcriptome profiles from high-throughput assays. Our integrative analysis of bulk WES and scRNA-seq from 32 human fibroblast cell lines reveals substantial phenotypic effects of somatic variation, including in healthy tissue.

Central to our approach is cardelino, a robust model for the probabilistic assignment of cells to clones based on variants contained in scRNA-seq reads. Our approach is conceptually related to de-multiplexing approaches for single-cell transcriptomes from multiple genetically distinct individuals (Kang *et al*., 2018). However, cardelino addresses a substantially more challenging problem: to distinguish cells from the same individual based on the typically small number of somatic variants (*e.g.* dozens) that segregate between clones in a population of cells.

Harnessing transcriptomic phenotypic information for cells assigned to clones in fibroblast lines, we identify substantial and convergent gene expression differences between clones across lines, which are notably enriched for pathways related to proliferation and the cell cycle. Analysis of clonal evolutionary dynamics using somatic variant allele frequency distributions from WES data reveals strong evidence for positive selection of clones in ten of 32 lines. These results support previous observations of clonal populations undergoing positive selection in normal human eyelid epidermis assayed by targeted DNA sequencing (Martincorena *et al*., 2015; Simons, 2016b; Martincorena *et al*., 2016; Simons, 2016a). We shed light on the phenotypic origin of this adaptive evolution, as we identify expression of gene sets implicated in proliferation and cancer such as the E2F and MYC pathways. This surprising result in healthy tissue, suggests pervasive inter-clonal phenotypic variation with important functional consequences.

While we apply cardelino to clonal trees from bulk WES data in this study, our method is general and can be used to assign cells to clonal trees inferred from either bulk or single-cell DNA-seq data. The methods presented here can be applied to any system in which somatic variants tag clonal populations of cells and can be accessed with scRNA-seq assays. Thus, cardelino will be applicable to high-resolution studies of clonal gene expression in both healthy and malignant cell populations as well as *in vitro* models.

Assignment of cells to clones relies on coverage of somatic variants in scRNA-seq reads, so cell populations with relatively fewer somatic variants may require full-length transcriptome sequencing at higher coverage per cell to enable confident assignments. Increasing both the number of genetically distinct individuals and the numbers of cells assayed per donor would improve the resolution at which differentially expressed genes and pathways can be identified. Larger sample sizes would also motivate further advances in the downstream analysis, for example to account for uncertainty in clonal tree inference and cell-clone assignment when assessing differentially expressed genes. Larger cell counts in particular would also enable joint modeling approaches combining DNA-seq and scRNA-seq data to infer clonal trees and simultaneously assign cells to clones. Our inference methods in cardelino are computationally efficient, so will comfortably scale to multi-site samples and many thousands of cells.

Taken together, our results highlight the utility of cardelino to study gene expression variability in clonal cell populations and suggest that even in nominally healthy human fibroblast cell lines there are clonal populations with growth advantages, opening new avenues to study cell behaviour in clonal populations.

## Methods

### The cardelino model

The input to cardelino is a clonal tree inferred using methods such as *Canopy*, yielding clonal fractions *F={f_1_,…, f_K_}*, where *f_k_* is the relative prevalence of a given clone 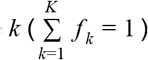, as well as a clonal tree configuration matrix *C* (an *N-* by-*K* binary matrix) for *N* variants and *K* clones, where *c _i,k,_=* 1 if somatic variant *i* is present in clone *k* and *c_i,k,_=* 0 otherwise. Given *C* and *F*, cardelino is aimed at assigning individual cells to one of *K* clones based on their expressed alleles using a probabilistic clustering model (see graphical model in **Supp. Fig. S17**). Based on scRNA-seq, we extract for each cell and variant that segregates between clones the number of sequencing reads that support the reference allele (reference read count) or the alternative allele (alternate read count) respectively. We denote the variant-by-cell matrix of alternate read counts by *A* and the variant-by-cell matrix of total read counts (sum of reference and alternate read counts) by *D*. Entries in *A* and *D* matrices are non-negative integers, with missing entries in the matrix *A* indicating zero read coverage for a given cell and variant.

Fundamentally, we model the alternate read count using a binomial model. For a given site in a given cell, there are two possibilities: the variant is “absent” in the clone the cell is assigned to or the variant is “present”, as encoded in the configuration matrix *C*. Thus, when considering the “success probability” *θ* for the binomial model, where success is defined as observing an alternate read in the scRNA-seq reads, we consider two (sets of) parameters: *θ_0_* for homozygous reference alleles (variant absent), and *Θ_1_* for heterozygous variants (variant present).

The prior probability that cell *j* belongs to clone *k* could be assigned as the clonal fraction *f_k’_*, but to avoid biasing cell assignment towards highly prevalent clones for cells with little read information (where an informative prior is particularly influential) we use a uniform prior such that *P(I_j_= k)* = 1/*K* for all *k*. Given this prior distribution, the posterior probability of cell *j* belonging to clone *k* can be expressed as:

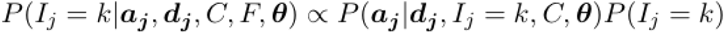

where *I_j_* is the identity of the specific clone cell *j* is assigned to, and ***a**_j_* and ***d**_j_* are the observed alternate and total read count vectors, respectively, for variants 1 to *N* in cell*j* The parameter vector *θ* is a set of the unknown binomial success parameters of binomial distributions for modelling the allelic read counts as described above. Specifically, *θ*_0_ denotes the binomial success rate for the alternative allele when *c_i,k,_*= 0 (variant absent), thereby accounting for sequencing errors or errors in the clonal tree configuration, and *θ*_1_={*θ*_1_,*θ*_2_…,*θ*_N_} denotes a vector of binomial parameters, one for each variant, for *c_i,k_*= 1. The latter binomial rates model the effect of allelic imbalance, which means the probability of observing alternate reads at frequencies that differ from 0.5 for true heterozygous sites (see **Supp.Methods** for details). The likelihood for cell *j* given an assignment to clone *k* follows then as a product of binomial distributions as follows,

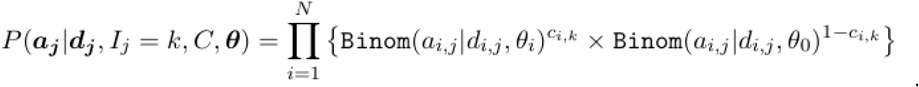

To capture the uncertainty in the binomial success probabilities, we introduce beta prior distributions on *θ*_0_ and **θ**_1_. To ensure sensible prior distributions, we estimate the beta parameters from the scRNA-seq at known germline heterozygous variants for highly expressed genes (**Supp. Fig. S18**). For example, in the fibroblast dataset considered here, this approach yielded prior parameters of beta(0.3, 29.7) for *θ_0_* and beta(2.25, 2.65) for *θ_i’_i*>0.

The posterior probability of cell assignment and the full posterior distribution of these parameters are inferred by a Gibbs sampler, and the details of the algorithm can be found in **Supp. Methods**, where we also present alternative simpler inference schemes using a constant Binomial success rates or an EM algorithm (see also **Supp. Fig. S1**). Despite the full Bayesian approach, cardelino is computationally efficient, enabling the assignment of hundreds of cells within minutes using a single compute node. These methods will comfortably scale to datasets with many thousands of cells.

### Cell culture

Dermal fibroblasts, derived from skin-punch samples from the shoulder of 32 donors (White British, age range 30-75), were obtained from the HipSci project (http://hipsci.org). Following thawing, fibroblasts were cultured in supplemented DMEM (high glucose, pyruvate, GlutaMAX (Life Technologies / 10569-010), with 10% FBS (Lab Tech / FB-1001) and 1% penicillin-streptomycin (Life Technologies / 15140122) added. 18 hours prior to collection, cells were trypsinised (Life Technologies / 25300054), counted, and seeded at a density of 100,000 cells per well (6 well plate).

### Cell pooling, capture and full-length transcript single-cell RNA sequencing

Cells were washed with PBS, trypsinised, and resuspended in PBS (Gibco / 14190-144) + 0.1% DAPI (AppliChem / A1001). Cells from three donors were pooled and consequently sorted on a Becton Dickinson INFLUX machine into plates containing 2uL/well lysis buffer. Single cells were sorted individually (using FSC-W vs FSC-H), and apoptotic cells were excluded using DAPI. Cells from each three-plex cell pool were sorted across four plates. Reverse transcription and cDNA amplification was performed according to the Smart-seq2 protocol (Picelli *et al*., 2014), and library preparation was performed using an Illumina Nextera kit. Samples were sequenced using paired-end 75bp reads on an Illumina HiSeq 2500 machine.

### Bulk whole-exome sequencing data and somatic variant calling

We obtained bulk whole-exome sequencing data from HipSci fibroblast (median read coverage: 254) and derived iPS cell lines (median read coverage: 79) released by the HipSci project (Streeter *et al*., 2016; Kilpinen *et al*., 2017). Sequenced reads were aligned to the GRCh37 build of the human reference genome (Church *et al*., 2011) using *bwa* (Li, 2013). To identify single-variant somatic mutation sites in the fibroblast lines, for each exome sample, we searched for sites with a non-reference base in the read pileup using *bcftools/mpileup* (Li *et al*., 2009). In the initial pre-filtering we retained sites with a per-sample coverage of at least 20 reads, at least three alternate reads in either fibroblast or iPS samples and an allele frequency less than 5% in the ExAC browser (Karczewski *et al*., 2017) and 1000 Genomes data (The 1000 Genomes Project Consortium, 2015). A Fisher exact test (Fisher, 1922) implemented in *bcftools/ad-bias* was then used to identify sites with significantly different variant allele frequency (VAF) in the exome data between fibroblast and iPS samples for a given line (Benjamini-Hochberg FDR < 10%). Sites were removed if any of the following conditions held: VAF < 1% or VAF > 45% in high-coverage fibroblast exome data; fewer than two reads supporting the alternative allele in the fibroblast sample; VAF > 80% in iPSC data (to filter out potential homozygous alternative mutations); neither the iPS VAF or fibroblast VAF was below 45% (to filter out variants with a “significant” difference in VAF but are more likely to be germline than somatic variants). We further filtered sites to require uniqueness of sites across donors as it is highly unlikely to observe the same point mutation in more than one individual, so such sites almost certainly represent technical artefacts.

### Identification of mutational signatures

Signature exposures were estimated using the *sigfit* package (Gori and Baez-Ortega, 2018), providing the COSMIC 30 signatures as reference (Forbes *et al*., 2017), and with a highest posterior density (HPD) threshold of 0.9. Signatures were determined to be significant when the HPD did not overlap zero. Two signatures (7 and 11) were significant in two or more donors.

### Identification of selection dynamics

Several methods have been developed to detect deviations from neutral growth in cell populations (Simons, 2016a; Williams *et al*., 2016, 2018; Martincorena *et al*., 2018). Methods such as dN/dS or models assessing the fit of neutral models to the data need a high number of mutations to determine selection/neutrality. Given the relatively low number of mutations found in the donors in this study, these models are not applicable. We used the package *SubClonalSelection* (https://github.com/marcjwilliams1/SubClonalSelection.jl) in *Julia 0.6.2* which works with a low number of mutations (> 100 mutations; Williams *et al*., 2018). The package simulates the fit of a neutral and a selection model to the allele frequency distribution, and returns a probability for the selection model to fit the data best.

At small allele frequencies the resolution of the allele frequency distribution is limited by the sequencing depth. We chose a conservative lower resolution limit of *f_min_* = 0.05 (Shin *et al*., 2017). At the upper end of the allele frequency distribution we chose a cut-off at *f_max_* =0.45 to account for ploidy (=2). For the classification of the donors, we introduced cut-offs on the resulting selection probability of the algorithm. Donors with a selection probability below 0.3 are classified as ‘neutral’, above 0.7 as ‘selected’. Donors which are neither ‘selected’ nor ‘neutral’ remain ‘undetermined’. See figure 2A and S3 for the results of the classification and fit of the models to the data.

### Single-cell gene expression quantification and quality control

Raw scRNA-seq data in CRAM format was converted to FASTQ format with *samtools* (v1.5), before reads were adapter-and quality-trimmed with *TrimGalore!* (github.com/FelixKrueger/TrimGalore) (Martin, 2011). We quantified transcript-level expression using Ensembl v75 transcripts (Flicek *et al*., 2014) by supplying trimmed reads to *Salmon* v0.8.2 and using the “--seqBias”, “--gcBias” and “VBOpt” options (Patro *et al*., 2017). Transcript-level expression values were summarised at gene level (estimated counts) and quality control of scRNA-seq data was done with the *scater* package (McCarthy *et al*., 2017) and normalisation with the *scran* package (Lun, Bach, *et al*., 2016; Lun, McCarthy, *et al*., 2016). Cells were retained for downstream analyses if they had at least 50,000 counts from endogenous genes, at least 5,000 genes with non-zero expression, less than 90% of counts from the 100 most-expressed genes in the cell, less than 20% of counts from ERCC spike-in sequences and a *Salmon* mapping rate of at least 40% (**Supp. Table S2**). This filtering approach retains 63.7% of assayed cells.

### Deconvolution of donors from pools

To increase experimental throughput in processing cells from multiple distinct donor individuals (*i.e.* lines), and to ensure an experimental design robust to batch effects, we pooled cells from three donors in each processing batch, as described above. As such, we do not know the donor identity of each cell at the time of sequencing and cell-donor identity must be inferred computationally. Thus, for both donor and, later, clone identity inference it is necessary to obtain the count of reads supporting the reference and alternative allele at informative germline and somatic variant sites. Trimmed FASTQ reads (described above) were aligned to the GRCh37 p13 genome with ERCC spike-in sequences with STAR in basic two-pass mode (Dobin *et al*., 2012) using the GENCODE v19 annotation with ERCC spike-in sequences (Searle *et al*., 2010). We further use *picard* (Broad Institute, 2015) and *GATK* version 3.8 (McKenna *et al*., 2010) to mark duplicate reads (*MarkDuplicates*), split cigar reads (*SplitNCigarReads*), realign indels (*IndelRealigner*), and recalibrate base scores (*BaseRecalibrator*).

For cell-donor assignment we used the *GATK HaplotypeCaller* to call variants from the processed single-cell BAM files at 304,405 biallelic SNP sites from dbSNP (Sherry *et al*., 2001) build 138 that are genotyped on the Illumina HumanCoreExome-12 chip, have MAF > 0.01, Hardy-Weinberg equilibrium P < 1e-03 and overlap protein-coding regions of the 1,000 most highly expressed genes in HipSci iPS cells (as determined from HipSci bulk RNA-seq data). We merged the per-cell VCF output from *GATK HaplotypeCaller* across all cells using *bcftools* version 1.7 (Danecek *et al*., 2011, 2016) and filtered variants to retain those with MAF > 0.01, quality score > 20 and read coverage in at least 3% of cells. We further filtered the variants to retain only those that featured in the set of variants in the high-quality, imputed, phased HipSci genotypes and filtered the HipSci donor genotype file to include the same set of variants.

We used the *donor_id* function in the cardelino package to assign cells to donors. This function assigns cells to donors by modelling alternative allele read counts with given genotypes of input donors. For a single germline variant, the three base genotypes (as minor allele counts) can be 0, 1 and 2. For doublet genotype profiles generated by combining pairs of donor genotypes, two additional combinatory genotypes, 0.5 and 1.5 are allowed. We assume that each genotype has a unique binomial distribution whose parameters are estimated by an EM algorithm in a framework similar to clone assignment (described above). When we enable doublet detection, the posterior probabilities that a cell comes from any of the donors provided, including doublet donors, are calculated for donor assignment. There are 490 available HipSci donors, so we run cardelino in two passes on each plate of scRNA-seq data separately. In the first pass, the model outputs the posterior probability that each cell belongs to one of the 490 HipSci donors, ignoring the possibility of doublets. In the second pass, only those donors with a posterior probability greater than 0.95 in at least one cell are considered by the model as possible donors and doublet detection is enabled. After the second pass, if the highest posterior probability is greater than 0.95, more than 25 variants have read coverage, and the doublet probability is less than 5% then we provisionally assign the cell to the donor with the highest posterior probability. If the provisionally assigned donor is one of the three donors known to have been pooled together for the specific plate, then we deem the cell to be confidently assigned to that donor, otherwise we deem the cell to have “unassigned” donor. With this approach, 97.4% of cells passing QC (see above) are confidently assigned to a donor (**Supp. Fig. S19**). Of the cells that are not confidently assigned to a donor, 2.1% are identified as doublets by cardelino and 0.5% remain “unassigned” due to low variant coverage or low posterior probability. Thus, we have 2,338 QC-passing, donor-assigned cells for clonal analysis.

### Clonal inference

We inferred the clonal structure of the fibroblast cell population for each of the 32 lines (donors) using Canopy (Jiang *et al*., 2016). We used read counts for the variant allele and total read counts at filtered somatic mutation sites from high-coverage whole-exome sequencing data from the fibroblast samples as input to Canopy. In addition to the variant filtering described above, input sites were further filtered for tree inference to those that had non-zero read coverage in at least one cell assigned to the corresponding line. We used the BIC model selection method in Canopy to choose the optimal number of clones per donor. Here, for each of the 32 lines, we considered the highest-likelihood clonal tree produced by Canopy, along with the estimated prevalence of each clone and the set of somatic variants tagging each clone as the given clonal tree for cell-clone assignment.

### Cell-clone assignment

For cell-clone assignment we required read the read counts supporting reference and alternative alleles at somatic variant sites. We used the *bcftools* version 1.7 *mpileup* and *call* methods to call variants at somatic variant sites derived from bulk whole-exome data, as described above, for all confidently assigned cells for each given line. Variant sites were filtered to retain variants with more than three reads observed across all cells for the line and quality greater than 20. We retained cells with at least two somatic variants with non-zero read coverage (2,044 cells across 32 lines). From the filtered VCF output of *bcftools* we obtained the number of reads supporting the alternative allele and the total read coverage for each somatic variant site with more than three reads covering the site, in total, across all the donor’s cells. In general, read coverage of somatic mutation sites in scRNA-seq is sparse, with over 80% of sites for a given cell have no overlapping reads. We used the scRNA-seq read counts at the line’s somatic variant sites to assign QC-passing cells from the line to clones using the *clone_id* function in the cardelino R package.

### Simulations to benchmark cell to clone assignment

We simulated data to test the performance of cardelino as follows. First, given a clonal tree configuration *C* (*N-*by-*K* binary matrix), a given number of cells are generated (e.g. 500, see below), whose genotypes are sampled from *K* clones following a multinomial distribution parameterised by clonal fractions *F*. Second, given a matrix *D* (*N*-by-*M* matrix) of sequencing coverage for *N* sites in *M* cells, we uniformly sample the coverage profiles from these *M* cells into a given number of cells for simulation. Third, after having the genotype *h=c,j* and the sequencing depth *d_y_* for variant *i* in cell *j* from the previous two steps, we can generate the read count *a _ij_* for the alternative allele by sampling from a binomial distribution with success parameter *θ_0_* if *h=0* or with an allele-specific expression parameter *θ* if *h=1*. Note, both *θ_0_* and *θ*_i_ are randomly generated from beta prior distributions, whose parameters are estimated from experimental data.

Based on the above simulation workflow, two simulation experiments are performed to evaluate the accuracy and robustness of cardelino. One simulation was performed with synthesizing 500 cells on each of 32 donors, where input parameters are from the observed matrices *C* and *D*, clonal fraction *F*, and cardelino-learned *θ* from each donor. This simulation tries to mimic all settings in each donor, which not only evaluates the accuracy of the model, but also reflects the quality of the data in each donor for clonal assignment. Alternatively, we change one of these parameters each time to systematically assess cardelino. The clonal configuration is defined by the number of clones, *K*, a perfect phylogenetic matrix *((K-* 1 )-by-*K)* including a base clone, and the number of unique variants per clonal branch *n*, which returns a configuration matrix *C* with a shape of *n(K-* 1 )-by-*K*. With setting *K* clones, one of all clonal tree structures are randomly selected to generate the clonal configuration matrix. Then the sequencing depth matrix *D* for these *n(K-* 1 ) variants are sampled from a donor with 439 variants across 151 cells (see distribution in **Supp. Fig. S20**). In order to increase or decrease the missingness rate of *D*, zero coverages are respectively added or removed linearly according to the expression level of the gene corresponding to the variant. The allelic expression balance can be adjusted by changing the parameters of its beta prior distribution. We set uniform clonal prevalence in the second simulation. With each parameter setting, 200 cells are randomly synthesized and this procedure is repeated 20 times to vary for the selection of variants and tree structure. When one setting parameter varies, others are used at the default values: number of mutations per clonal branch = 10, variant coverage = 0.2, clone number = 4, mean fraction of alternative alleles (i.e. mean ö_1_) = 0.44, variance of allelic imbalance (i.e. 1/(shape1+shape2) of beta prior) = 0.21.

### Differential expression, pathway and variance component analysis

Expression analyses between clones required further filtering of cells for each line. Analyses were conducted using cells that passed the following filtering procedure for each line: (1) clones identified in the line were retained if at least three cells were confidently assigned to the clone; (2) cells were retained if they were confidently assigned to a retained clone. Lines were retained for DE testing if they had at least 15 cells assigned to retained clones, allowing us to conduct expression analyses for 31 out of the 32 lines (all except *vils*).

Differential gene expression (DE) testing was conducted using the quasi-likelihood F-test method (Lund *et al*., 2012) in the *edgeR* package (Robinson *et al*., 2010; McCarthy *et al*., 2012) as recommended by Soneson and Robinson (Soneson and Robinson, 2018). To test for differences in expression between cells assigned to different clones in a donor, we fit a linear model for single-cell gene expression with cellular detection rate (proportion of QC-filtered genes expressed in a cell; numeric value), plate on which the cell was processed (a factor) and assigned clone (a factor) as predictor variables. The quasi-likelihood F test was used to identify genes with: (1) any difference in average expression level between clones (analogous to analysis of variance), and (2) differences in average expression between all pairs of clones (“pairwise contrasts”). We considered 10,876 genes that were sufficiently expressed (an average count >1 across cells in all donors) to test for differential expression.

To test for significance of overlap of DE genes across donors, we sampled sets of genes without replacement the same size as the number of DE genes (FDR < 10%) for each line. For each permutation set, we then computed the number of sampled genes shared between between donors. We repeated this procedure 1,000 times to obtain distributions for the number of DE genes shared by multiple donors if shared genes were obtained purely by chance.

Gene set enrichment (pathway) analyses were conducted using the *camera* (Wu and Smyth, 2012) method in the *limma* package (Smyth, 2004; Ritchie *et al*., 2015). Using log_2_-fold-change test statistics for 10,876 genes for pairwise contrasts between clones from the *edgeR* models above as input, we applied *camera* to test for enrichment for the 50 Hallmark gene sets from MSigDB, the Molecular Signatures Database (Liberzon *et al*., 2011). For all differential expression and pathway analyses we adjusted for multiple testing by estimating the false discovery rate (FDR) using independent hypothesis weighting (Ignatiadis *et al*., 2016), as implemented in the *IHW* package, with average gene expression supplied as the independent covariate.

Expression variance across cells is decomposed into multiple components in a linear mixed model, including cellular detection rate (proportion of genes with non-zero expression per cell) as a fixed effect and plate (i.e. experimental batch), donor (i.e. line; only when combining cells across all donors) and clone (nested within donor for combined-donor analysis) as random effects. We fit the linear mixed model on a per-gene basis using the *variancePartition* R package (Hoffman and Schadt, 2016).

### Code availability

The cardelino methods are implemented in an open-source, publicly available R package (github.com/PMBio/cardelino). The code used to process and analyse the data is available (github.com/davismcc/fibroblast-clonality), with a reproducible workflow implemented in Snakemake (Köster and Rahmann, 2012). Descriptions of how to reproduce the data processing and analysis workflows, with html output showing code and all figures presented in this paper, are available at davismcc.github.io/fibroblast-clonality. Docker images providing the computing environment and software used for data processing (hub.docker.com/r/davismcc/fibroblast-clonality/) and data analyses in R (hub.docker.com/r/davismcc/r-singlecell-img/) are publicly available.

### Data availability

Single-cell RNA-seq data have been deposited in the ArrayExpress database at EMBL-EBI (www.ebi.ac.uk/arrayexpress) under accession number E-MTAB-7167. Whole-exome sequencing data is available through the HipSci portal (www.hipsci.org). Metadata, processed data and large results files are available under the DOI 10.5281/zenodo.1403510 (doi.org/10.5281/zenodo.1403510).

## Supporting information

## Author contributions

R.R., T.H. and S.A.T. conceived and planned the experiments. R.R. and T.H. carried out the experiments. Y.H., D.J.M. and O.S. developed the computational methods. Y.H. and D.J.M. wrote the software. Y.H. planned and carried out the simulations. The HipSci Consortium provided the cell lines and exome sequencing data. P.D. conducted somatic variant calling from exome sequencing data. D.J.G. advised on somatic variant calling approaches and the mutational signatures analysis carried out by R.R. D.J.M. and M.J.B. developed data processing workflows and D.J.M. processed the single-cell RNA-sequencing data. D.J.K. conducted the selection analyses, supervised by B.D.S. D.J.M. and Y.H. carried out clonal inference and cell assignment analyses. D.J.M. conducted differential gene and pathway expression analyses and integrated the computational analyses into a reproducible workflow. D.J.M. and R.R. took the lead in writing the manuscript. D.J.M., R.R. and Y.H. drafted the manuscript and designed the figures. W.W. suggested improvements to somatic variant calling and differential expression analyses. S.A.T. and O.S. conceived of the study, planned and supervised the work. All authors contributed to the interpretation of results and commented on and approved the final manuscript. The HipSci Consortium generated and provided early access to the fibroblast lines used in this work (see **Supp. Material** for a full list of consortium members).

## Acknowledgements

We would like to thank David Jörg for highly constructive discussions. We would like to acknowledge the Wellcome Sanger Institute Cellular Genetics and Phenotyping teams (in particular, Alex Alderton, Celine Gomez, Rachel Boyd, Sharad Patel and Sam Barnett) and DNA pipelines for their invaluable assistance in generating the data for this study.

