## Supplementary material for "Cardelino: Integrating whole exomes and single-cell transcriptomes to reveal phenotypic impact of somatic variants"

*[1]European Molecular Biology Laboratory, European Bioinformatics Institute, Wellcome Genome Campus, CB10 1SD Hinxton, Cambridge, UK; [2]Wellcome Sanger Institute, Wellcome Genome Campus, Hinxton, CB10 1SA, UK; [3]European Molecular Biology Laboratory, Genome Biology Unit, 69117 Heidelberg, Germany; [4]St Vincent's Institute of Medical Research, Fitzroy, Victoria 3065, Australia. [5]Cavendish Laboratory, Department of Physics, JJ Thomson Avenue, Cambridge, CB3 0HE, UK. [6]The Wellcome Trust/Cancer Research UK Gurdon Institute, University of Cambridge, Cambridge, CB2 1QN, UK. [7]The Wellcome Trust/Medical Research Council Stem Cell Institute, University of Cambridge, Cambridge, UK. Department of Bioinformatics and Computational Biology, The University of Texas MD Anderson Cancer Center, Houston, Texas 77030, USA. [9]Division of Computational Genomics and Systems Genetics, German Cancer Research Center (DKFZ), 69120, Heidelberg, Germany.*

** These authors contributed equally to this work.*

*# Corresponding authors.*

ORCIDs:

- DJM: 0000-0002-2218-6833
- RR: 0000-0002-4453-3357
- YH: 0000-0003-3124-9186
- DJK: 0000-0003-3597-6591
- PD:
- MJB: 0000-0002-8431-3180
- TH:
- WW: 0000-0003-0617-9438
- DJG: 0000-0002-1529-1862
- BDS: 0000-0002-3875-7071
- OS: 0000-0002-8818-7193
- ST: 0000-0002-6294-6366

### HipSci consortium members

Helena Kilpinen[2,8], Angela Goncalves[2], Andreas Leha[2,10], Vackar Afzal[3], Kaur Alasoo[2], Sofie Ashford[4], Sendu Bala[2], Dalila Bensaddek[3], Marc Jan Bonder[1], Francesco Paolo Casale[1], Oliver J Culley[5], Anna Cuomo[1], Petr Danecek[2], Adam Faulconbridge[1], Peter W Harrison[1], Annie Kathuria[5] , Davis J McCarthy[1,9], Shane A McCarthy[2], Ruta Meleckyte[5], Yasin Memari[2], Bogdan Mirauta[1], Nathalie Moens[5], Filipa Soares[6], Alice Mann[2], Daniel Seaton[1], Ian Streeter[1], Chukwuma A Agu[2], Alex Alderton[2], Rachel Nelson[2], Sarah Harper[2], Minal Patel[2], Alistair White[2], Sharad R Patel[2], Laura Clarke[1], Reena Halai[2], Christopher M Kirton[2], Anja KolbKokocinski[2], Philip Beales[8], Ewan Birney[1], Davide Danovi[5], Angus I Lamond[3], Willem H Ouwehand[2,4,7], Ludovic Vallier[2,6], Fiona M Watt[5], Richard Durbin[2,11], Oliver Stegle[1,12,13] , Daniel J Gaffney[2]

[1]European Molecular Biology Laboratory, European Bioinformatics Institute, Wellcome Genome Campus, Hinxton, Cambridge, CB10 1SD, United Kingdom.
[2]Wellcome Trust Sanger Institute, Wellcome Genome Campus, Hinxton, Cambridge, CB10 1SA, United Kingdom.
[3]Centre for Gene Regulation & Expression, School of Life Sciences, University of Dundee, DD1 5EH, United Kingdom.
[4]Department of Haematology, University of Cambridge, Cambridge, United Kingdom.
[5]Centre for Stem Cells & Regenerative Medicine, King's College London, Tower Wing, Guy's Hospital, Great Maze Pond, London SE1 9RT, United Kingdom.
[6]Wellcome Trust and MRC Cambridge Stem Cell Institute and Biomedical Research Centre, Anne McLaren Laboratory, University of Cambridge, CB2 0SZ, United Kingdom.
[7]NHS Blood and Transplant, Cambridge Biomedical Campus, Cambridge, United Kingdom.
[8]UCL Great Ormond Street Institute of Child Health, University College London, London WC1N 1EH, United Kingdom.
[9]St Vincent's Institute of Medical Research, Fitzroy Victoria 3065, Australia.
[10]Department of Medical Statistics, University Medical Center Göttingen, Humboldtallee 32, 37073 Göttingen, Germany.
[11]Department of Genetics, University of Cambridge, Cambridge, United Kingdom.
[12]European Molecular Biology Laboratory, Genome Biology Unit, 69117 Heidelberg, Germany.
[13]Division of Computational Genomics and Systems Genetics, German Cancer Research Center (DKFZ), 69120, Heidelberg, Germany.

| Line name | Gender | Age | Number of Variants | Signature 7 Mean Exposure | Number Clones With Cells | Minimum Hamming Distance | Total cells | Assigned Cells | Proportion Assigned Cells |
|---|---|---|---|---|---|---|---|---|---|
| euts | male | 60-64 | 292 | 0.585 | 3 | 29 | 79 | 78 | 0.987 |
| fawm | female | 70-74 | 101 | 0.337 | 3 | 5 | 53 | 47 | 0.887 |
| feec | male | 60-64 | 170 | 0.281 | 4 | 5 | 75 | 64 | 0.853 |
| fikt | male | 50-54 | 142 | 0.378 | 3 | 13 | 39 | 36 | 0.923 |
| garx | female | 50-54 | 592 | 0.670 | 3 | 57 | 70 | 69 | 0.986 |
| gesg | male | 60-64 | 157 | 0.372 | 3 | 23 | 105 | 101 | 0.962 |
| heja | male | 70-74 | 192 | 0.266 | 3 | 16 | 50 | 50 | 1.000 |
| hipn | male | 55-59 | 59 | 0.019 | 3 | 8 | 62 | 49 | 0.790 |
| ieki | female | 55-59 | 82 | 0.381 | 3 | 7 | 58 | 26 | 0.448 |
| joxm | female | 45-49 | 612 | 0.609 | 3 | 41 | 79 | 77 | 0.975 |
| kuco | female | 65-69 | 41 | 0.112 | 2 | 9 | 48 | 48 | 1.000 |
| laey | female | 70-74 | 278 | 0.532 | 3 | 36 | 55 | 55 | 1.000 |
| lexy | female | 60-64 | 55 | 0.069 | 3 | 6 | 63 | 63 | 1.000 |
| naju | male | 60-64 | 85 | 0.296 | 2 | 13 | 44 | 44 | 1.000 |
| nusw | male | 65-69 | 62 | 0.091 | 3 | 3 | 60 | 20 | 0.333 |
| oaaz | male | 70-74 | 90 | 0.172 | 3 | 17 | 38 | 37 | 0.974 |
| oilg | male | 65-69 | 211 | 0.505 | 3 | 2 | 90 | 57 | 0.633 |
| pipw | male | 50-54 | 233 | 0.551 | 3 | 34 | 107 | 107 | 1.000 |
| puie | male | 60-64 | 117 | 0.448 | 3 | 10 | 41 | 41 | 1.000 |
| qayj | female | 60-64 | 46 | 0.035 | 3 | 7 | 97 | 59 | 0.608 |
| qolg | male | 35-39 | 120 | 0.381 | 2 | 23 | 36 | 36 | 1.000 |
| qonc | female | 65-69 | 131 | 0.406 | 3 | 7 | 58 | 43 | 0.741 |
| rozh | female | 65-69 | 79 | 0.173 | 4 | 2 | 91 | 42 | 0.462 |
| sehl | female | 55-59 | 178 | 0.527 | 4 | 2 | 30 | 24 | 0.800 |
| ualf | female | 55-59 | 325 | 0.540 | 3 | 29 | 89 | 88 | 0.989 |
| vass | female | 30-34 | 412 | 0.647 | 3 | 35 | 37 | 37 | 1.000 |
| vils | female | 35-39 | 51 | 0.206 | 4 | 1 | 37 | 4 | 0.108 |
| vuna | female | 65-69 | 135 | 0.456 | 2 | 33 | 71 | 71 | 1.000 |
| wahn | female | 65-69 | 496 | 0.605 | 3 | 52 | 82 | 77 | 0.939 |
| wetu | female | 55-59 | 73 | 0.212 | 3 | 8 | 77 | 66 | 0.857 |
| xugn | male | 65-69 | 124 | 0.398 | 3 | 8 | 35 | 34 | 0.971 |
| zoxy | female | 60-64 | 61 | 0.117 | 3 | 8 | 88 | 82 | 0.932 |

**Table S1:** Biological and technical metadata for each of the 32 HipSci human fibroblast lines used. Number of variants refers to somatic variants identified from whole-exome sequencing data (**Methods**); Signature 7 exposure refers to Signature 7 (UV) from the COSMIC set of mutational signatures; Minimum Hamming distance denotes the minimum number of variants distinguishing between two clones in the inferred clonal tree for the line (**Methods**).

| | Metric | Min. | 1st Qu. | Median | Mean | 3rd Qu. | Max. | % passing filter |
|---|---|---|---|---|---|---|---|---|
| **Before QC filtering** | Total counts from endog. genes | 178 | 123,489 | 383,929 | 442,353 | 621,738 | 5,833,292 | - |
| | Total genes expressed | 174 | 6,772 | 10,446 | 8,801 | 11,790 | 16,243 | - |
| | % counts from ERCCs | 0 | 0.97 | 1.81 | 14.47 | 3.34 | 99.90 | - |
| | % counts top 100 expressed genes | 29.4 | 40.8 | 55.6 | 57.8 | 62.8 | 100.0 | - |
| | % reads mapped | 7.69 | 68.71 | 75.59 | 74.80 | 81.67 | 100.0 | - |
| **After QC filtering** | Total counts from endog. genes | 50,464 | 316,033 | 484,887 | 559,742 | 710,028 | 2,659,889 | 80.6 |
| | Total genes expressed | 5,083 | 9,960 | 11,108 | 10,846 | 12,100 | 14,804 | 79.3 |
| | % counts from ERCCs | 0.001 | 0.96 | 1.63 | 1.86 | 2.39 | 18.1 | 85.3 |
| | % counts top 100 expressed genes | 29.4 | 38.6 | 52.4 | 49.2 | 58.2 | 89.0 | 86.1 |
| | % reads mapped | 44.1 | 70.3 | 76.0 | 74.8 | 79.1 | 92.7 | 99.3 |

**Table S2:** Summaries of QC metrics for single-cell RNA-seq data before and after QC filtering. Cells were required to have more than 50,000 counts from endogenous genes, more than 5,000 genes expressed (*i.e*. with non-zero expression), less than 20% of counts from ERCC transcripts, less than 90% of counts from the 100 most-expressed genes in the cell and at least 40% of reads mapped using *Salmon*. Metrics were computed using the *scater* package (**Methods**).

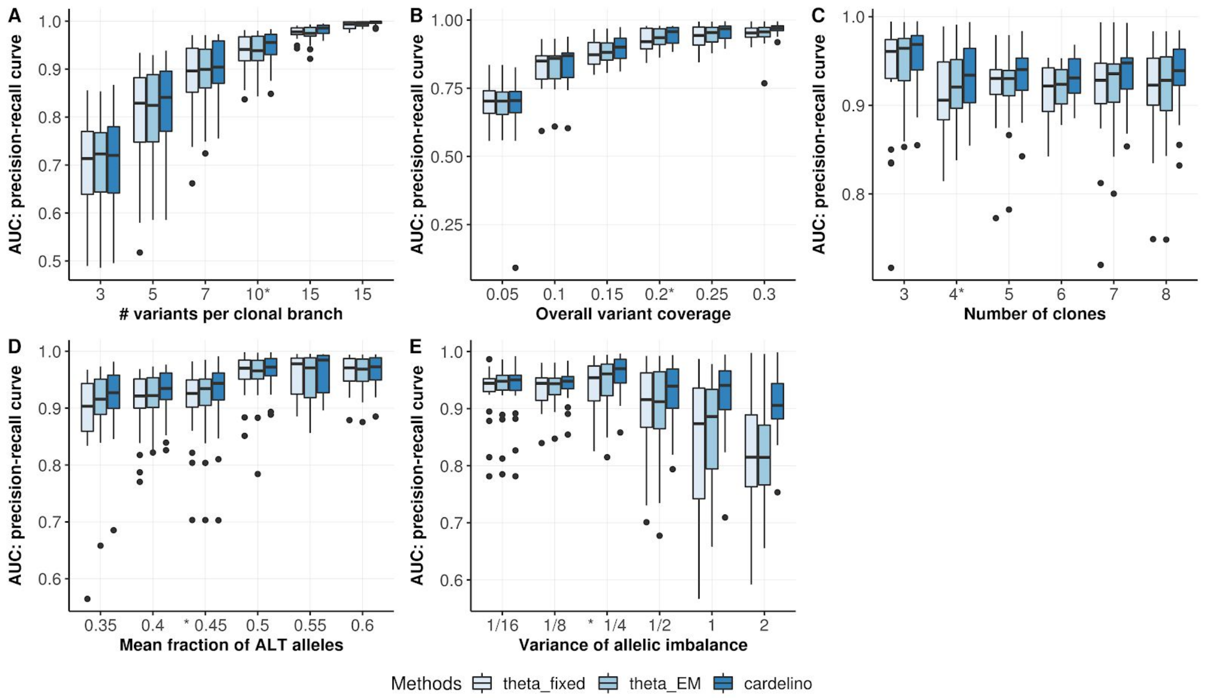


**Fig S1.** Assessment of cell assignment to clones, considering three variants of the cardelino model across a variety of simulation settings. Underlying all models is the approach of modeling read counts supporting the alternative allele at a site of somatic variation using a binomial model. For a given site in a given cell, there are two possibilities: the variant is "absent" in the cell (*i.e.* the cell has the homozygous reference genotype at that position) or the variant is "present" (*i.e.* the cell is heterozygous at that position). Thus, when considering the "success probability" $\boldsymbol{\theta}$ for the binomial model, where here success is defined as observing a read supporting the alternative allele for a variant, we have two (sets of) parameters with different values that need to be estimated: $\theta_0$ for homozygous reference alleles (variant absent), and $\boldsymbol{\theta}_1$ for the case with heterozygous variants (variant present). The three methods considered here are all based on the same clone-mixture model with a beta-binomial error model as in cardelino (**Methods**), but employ different strategies for estimating the binomial parameters $\theta_0$ and $\theta_1$ in the mixture model. Method **theta_fixed** considers *a priori defined* parameters ($\theta_0$=0.01 and $\boldsymbol{\theta}_1$=0.5), which corresponds to the assumption of a low, fixed sequencing error probability ($\theta_0$) and that from a heterozygous site (variant present) we are as likely to see a reference read as an alternative read ($\boldsymbol{\theta}_1$=0.5) and that these values are constant across all variants. The **theta_EM** model extends this base model by using an EM algorithm to obtain global maximum-likelihood estimates of $\theta_0$ and $\boldsymbol{\theta}_1$, again assuming the same error model across all variants. Finally, **cardelino** considers a vector $\boldsymbol{\theta}_1$ of variant-specific (same as gene-specific) parameters to account for gene-specific differences in allelic imbalance in gene expression. The cardelino model is therefore more flexible and able to model greater variation in read count distributions between variants due to either biological or technical factors. Cardelino uses a Gibbs sampler for parameter inference (see **Methods**). These three methods were assessed using simulated data, varying (**A**) the number of informative variants per clonal branch, (**B**) the average fraction of clone-specific variants detectable in scRNA-seq reads per cell, (**C**) the total number of clones, (**D**) the mean fraction of alternative allelic reads, i.e., mean $\boldsymbol{\theta}_1$ over genes (default 0.44), (**E**) the variance of allelic imbalance across genes, i.e., 1/(shape1+shape2), when sampling these values from a beta prior distribution (default 0.21). Default parameter values are marked with an asterisk and are retained when varything other parameters. In particular, when simulating gene to gene variability in allelic imbalance, the full cardelino model significantly outperforms alternative methods.

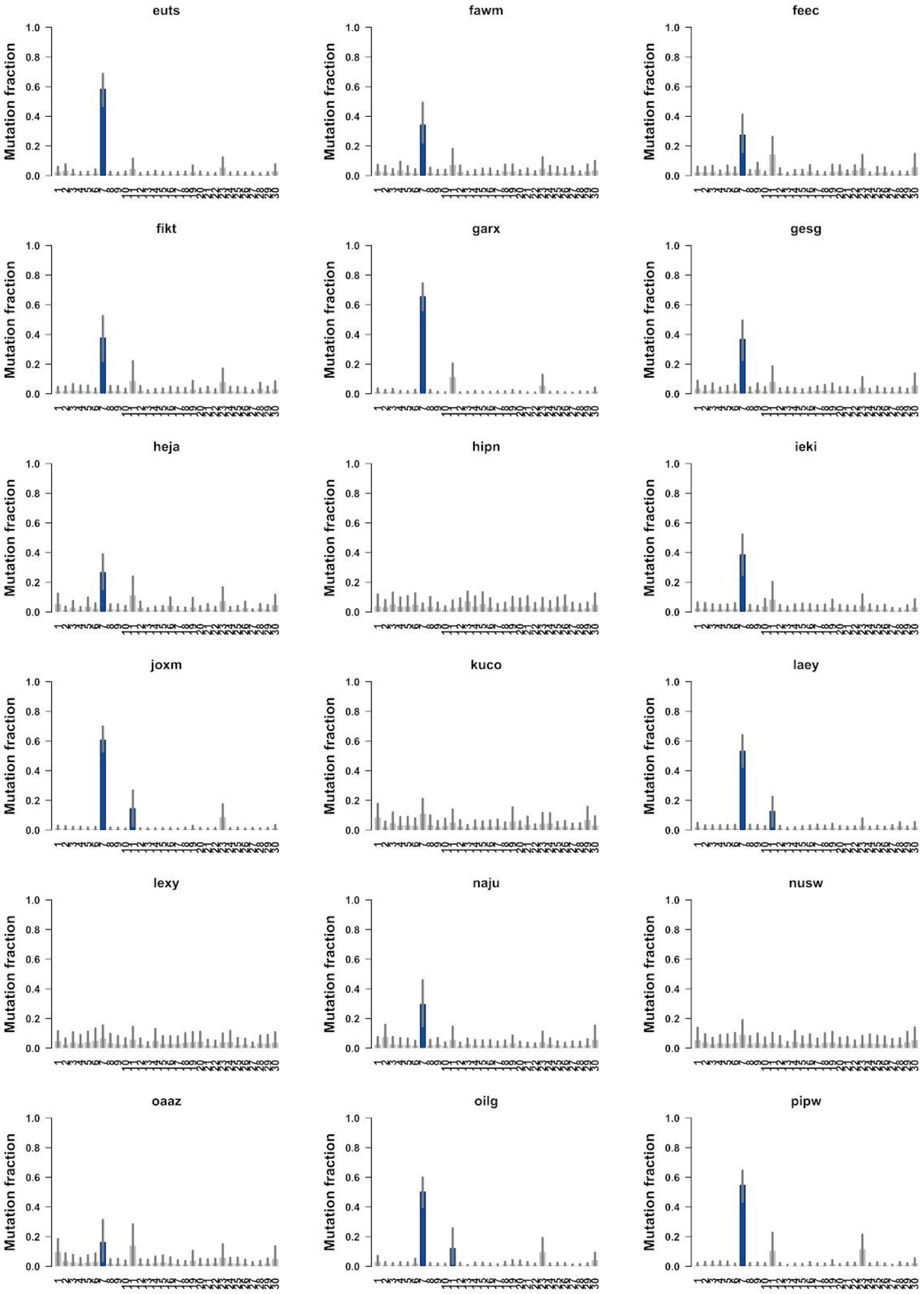
euts
fawm
feec
fikt
garx
gesg
heja
hipn
ieki
joxm
kuco
laey
lexy
naju
nusw
oaaz
oilg
pipw
Mutation fraction
0.0
0.2
0.4
0.6
0.8
1.0

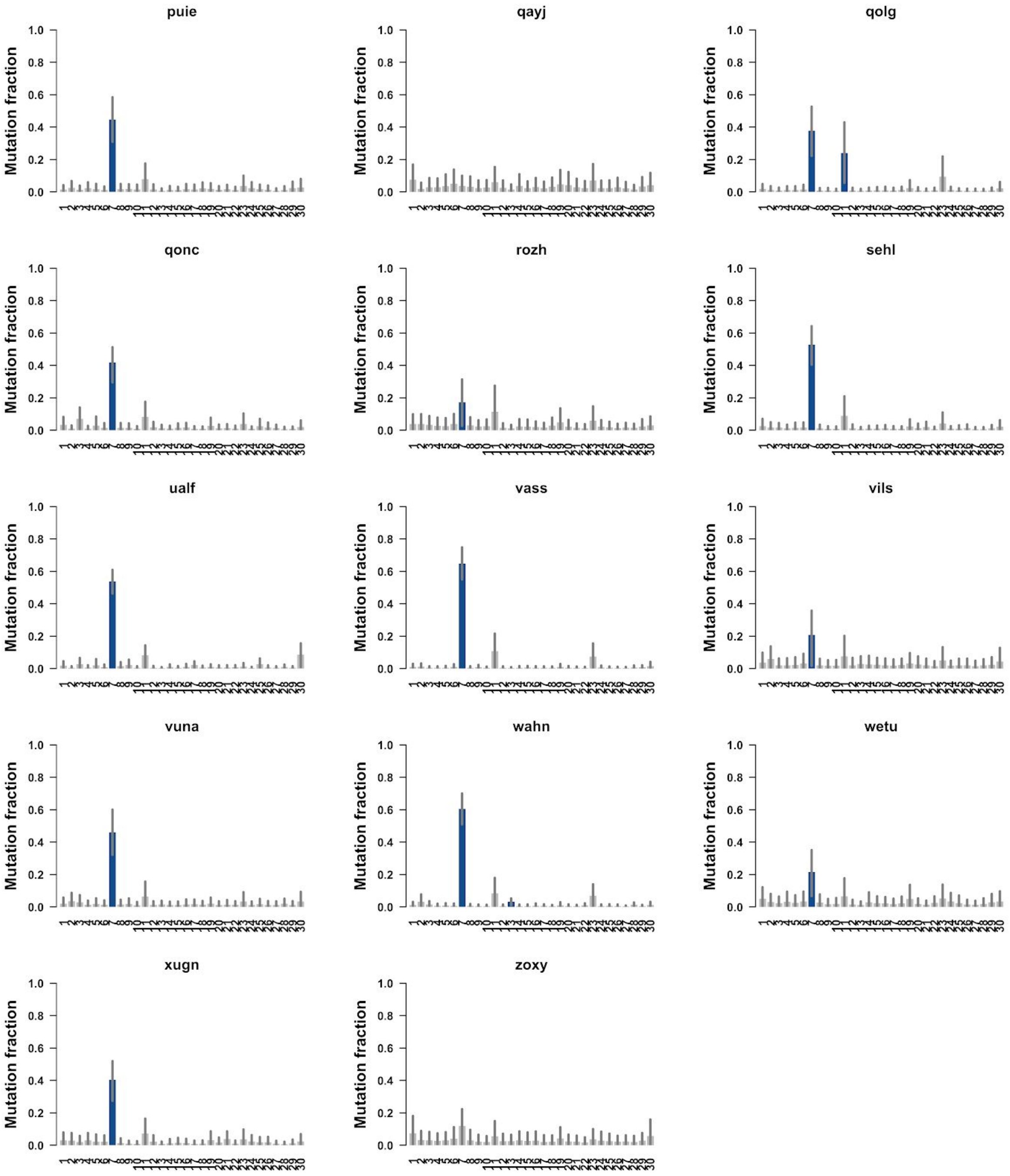


**Fig S2:** Estimated mutational signature exposures based upon the tri-nucleotide context of somatic SNVs called from whole-exome sequencing (WES) data for 32 HipSci human fibroblast lines. The x-axis shows 30 COSMIC mutational signatures, in order, and the y-axis shows estimated exposures (mutation fraction) using the *sigfit* package (**Methods**), with significant signatures highlighted in blue. Across lines, the only significant signatures are Signature 7 (UV mutagenic process) and Signature 11.

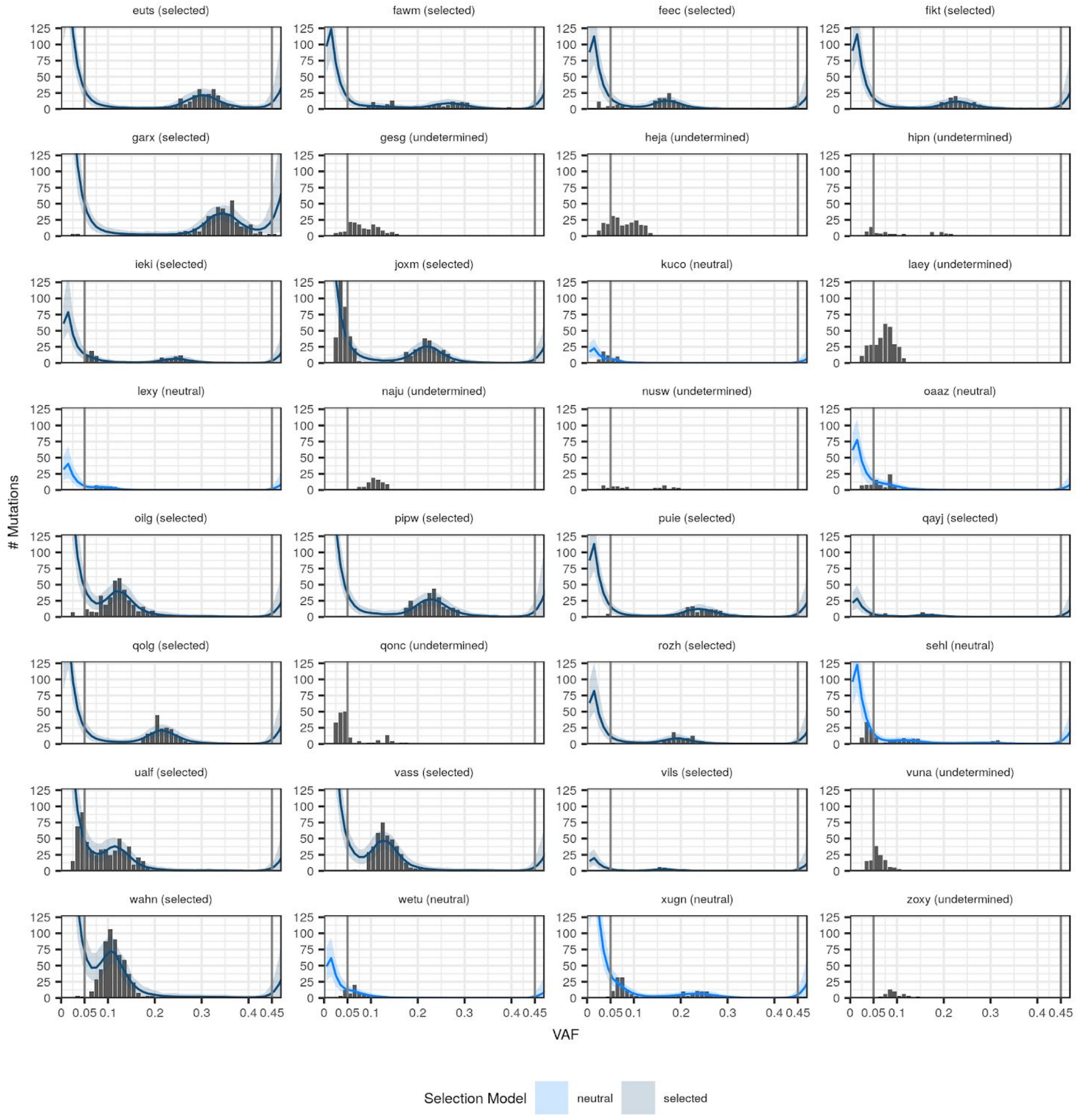


**Fig S3**: Allele frequency distributions for somatic variants called from WES data for the 32 fibroblast lines. The grey lines indicate the cut-offs on the allele frequency distribution (**Methods**). The blue lines are the fits of the neutral/selected model (shading 95% confidence interval).

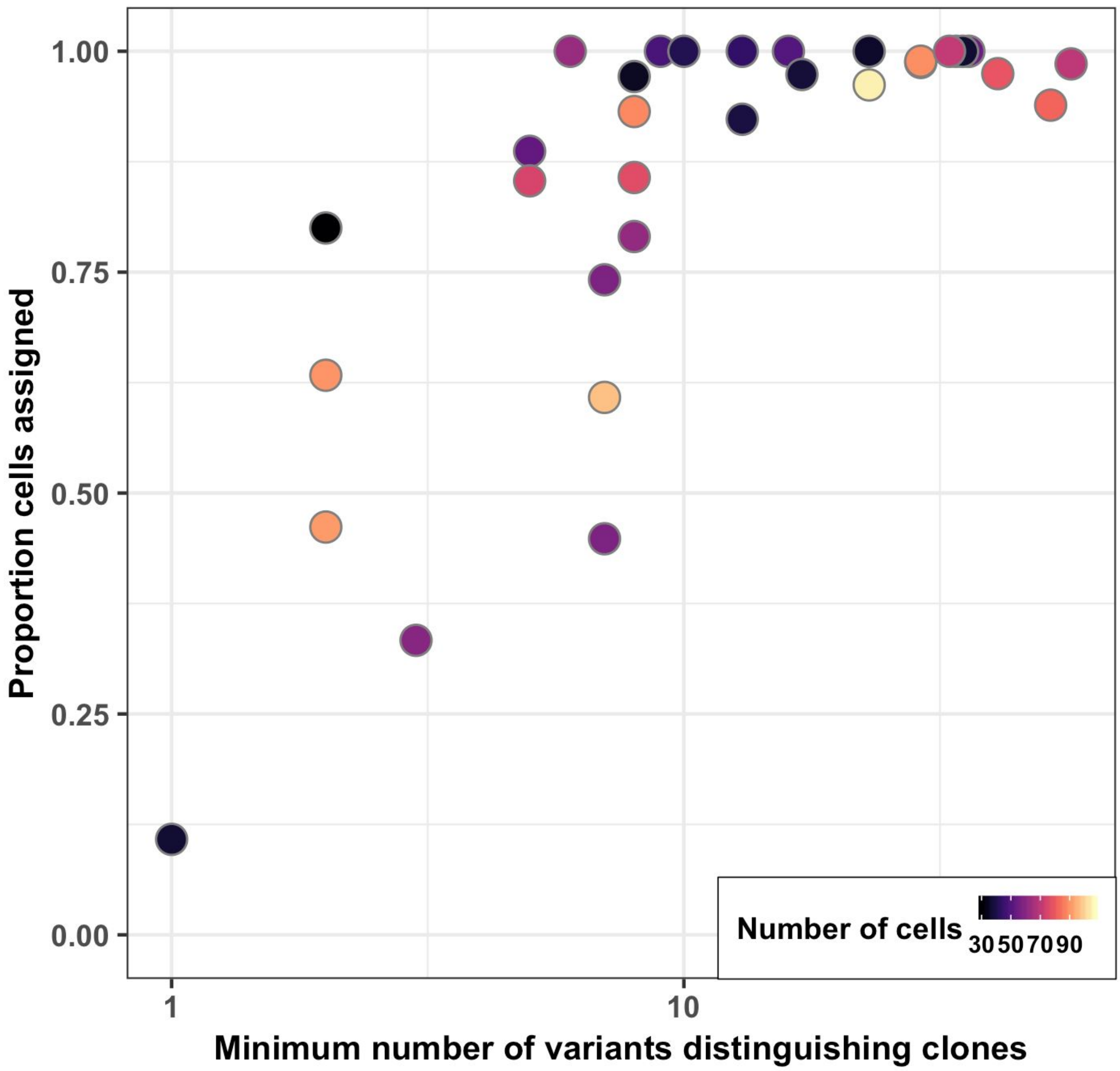


**Fig. S4:** Scatter plot of the fraction of cells assigned in each cell line using cardelino (at posterior probability > 0.5) as a function of the minimum number of clone-specific variants for the corresponding line (minimum Hamming distance between clones for a given donor), for 32 fibroblast lines. Total number of cells that were considered for this analysis (QC passed) per line indicated by colour.

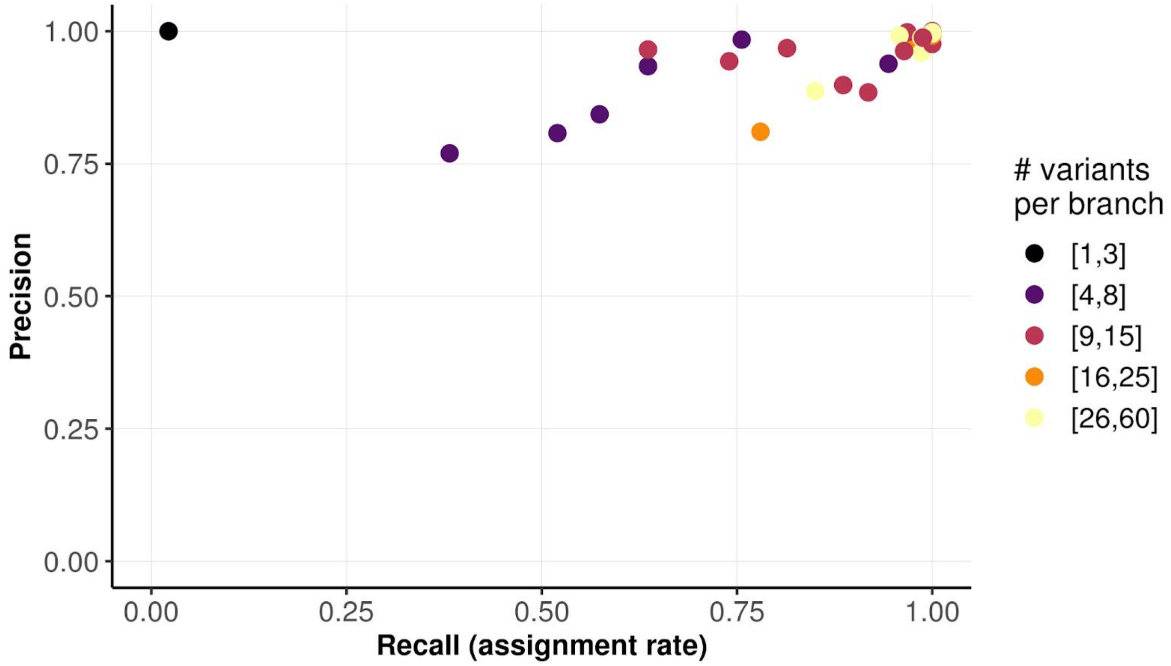


**Fig S5**. Scatter plot of recall (assignment rate) versus precision (assignment accuracy) when assigning cells using cardelino ( (at posterior probability > 0.5). Shown are data from for 32 simulated lines, using parameters that match the observed data characteristics in the set of 32 real fibroblast lines (**Methods**). The average number of variants per clonal branch (*i.e.*, #variant / (#clone - 1)) is shown by point colour (slightly different from Fig S4 which uses the minimum number of variants distinguishing between pairs of clones, as shown in Fig3a). Lines with fewer informative variants per branch tend to have lower assignment rates, but the precision remains high.

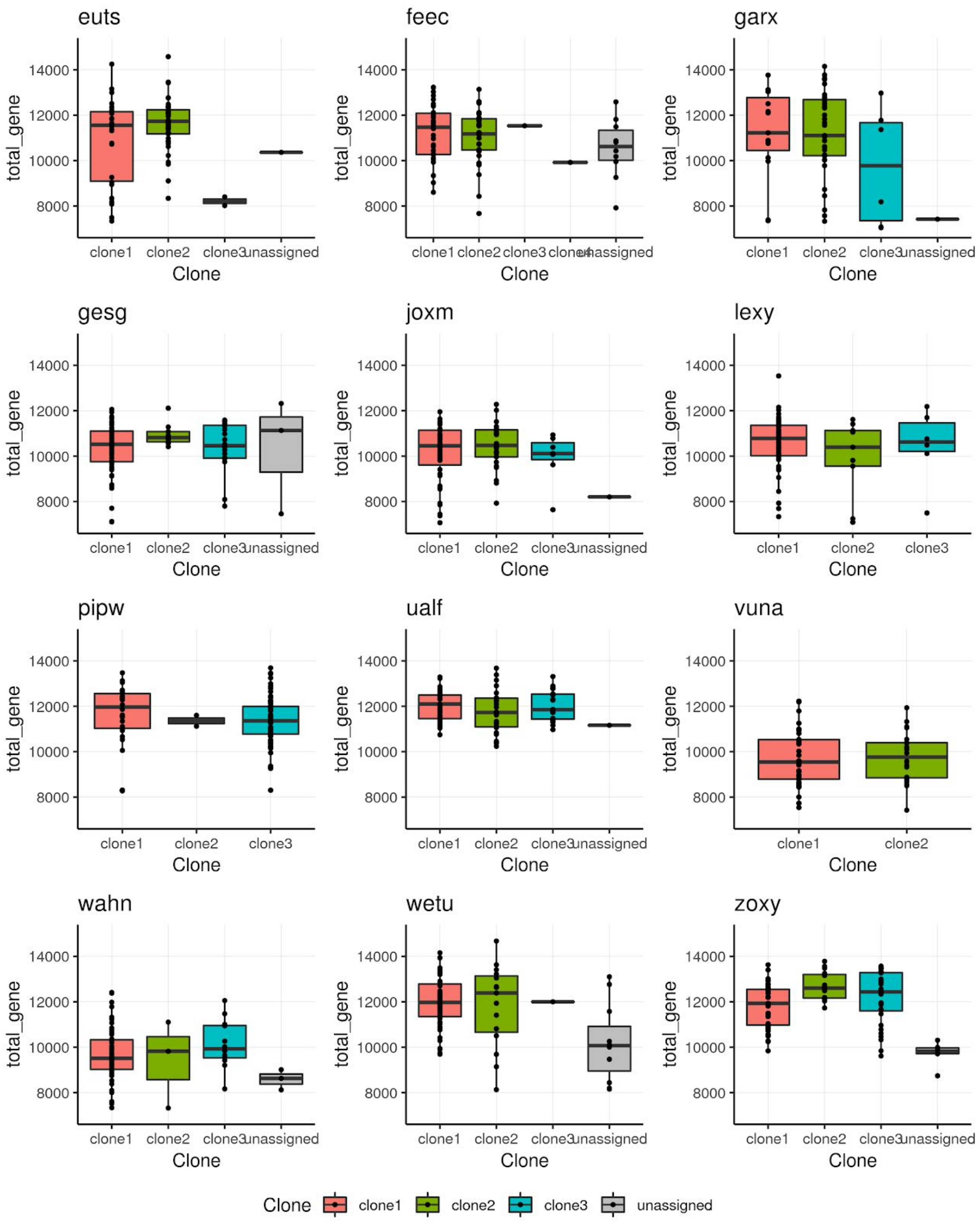


**Fig S6**: Boxplots of the total number of expressed genes in each cell, grouped by the clone assigned by *cardelino*. Twelve lines with more than 60 assignable cells are presented. Globally, clone assignment is not linked to the total number of expressed genes in a given cell.

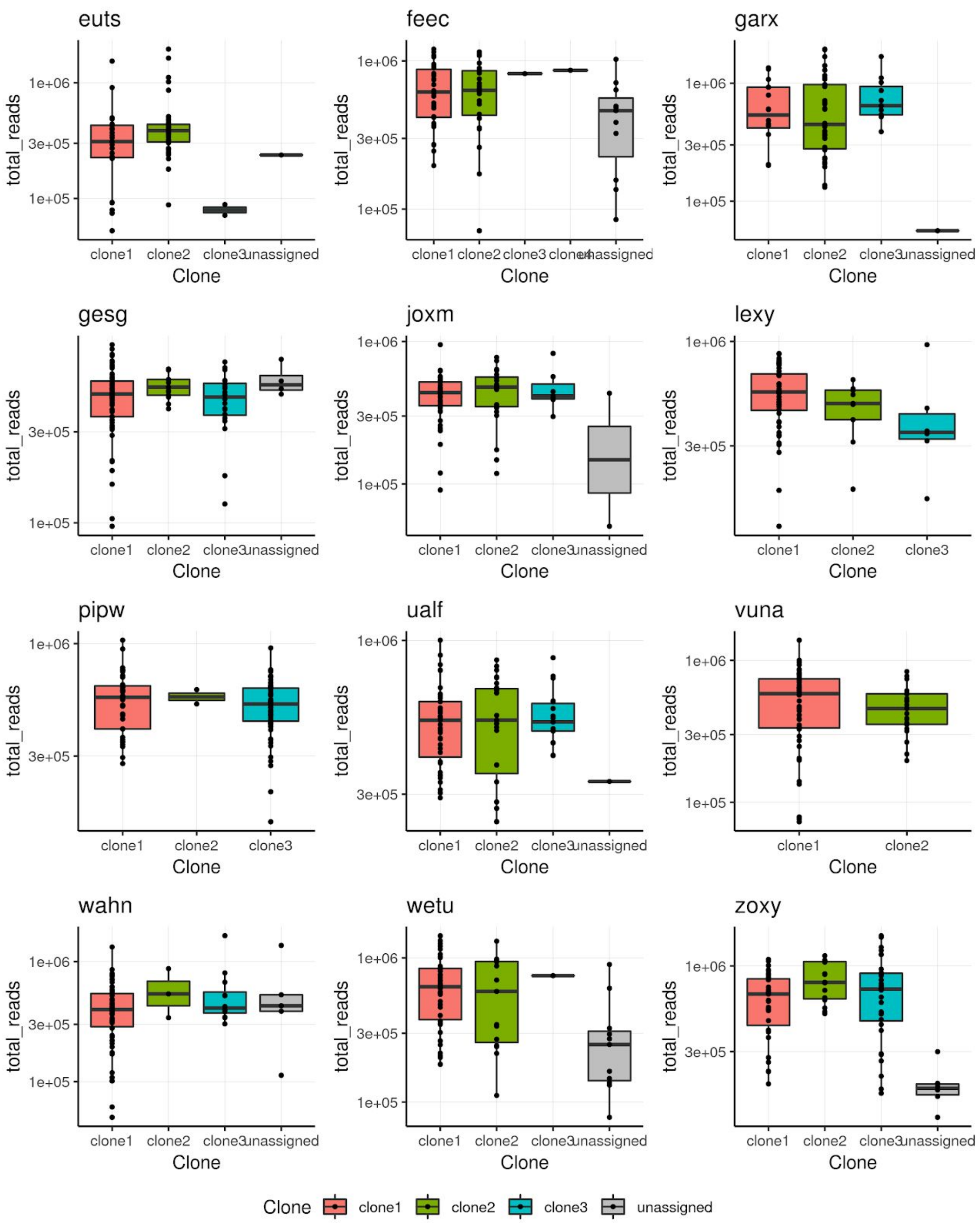


**Fig S7**: Boxplots of the total number of sequenced read counts from endogenous genes in each cell, grouped by the clone assigned by *cardelino*. Twelve lines with more than 60 assignable cells are presented. Globally, clone assignment is not linked to the total number of read counts in a given cell.

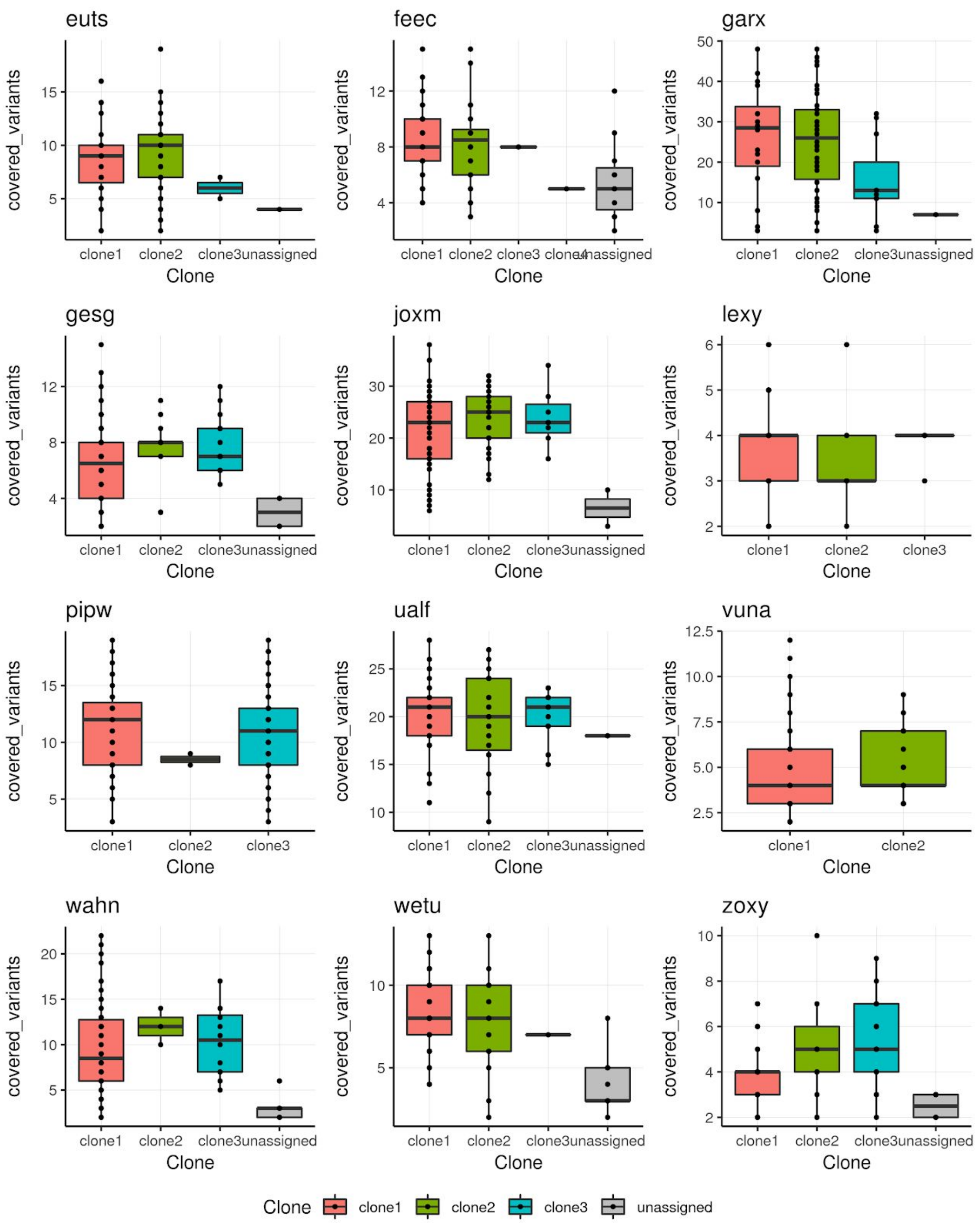


**Fig S8**: Boxplots of the number of variants for clone identification with read coverage in each cell, grouped by the clone assigned by *cardelino*. Twelve lines with more than 60 assignable cells are presented. Globally, clone assignment is not linked to the number expressed variant loci in a given cell.

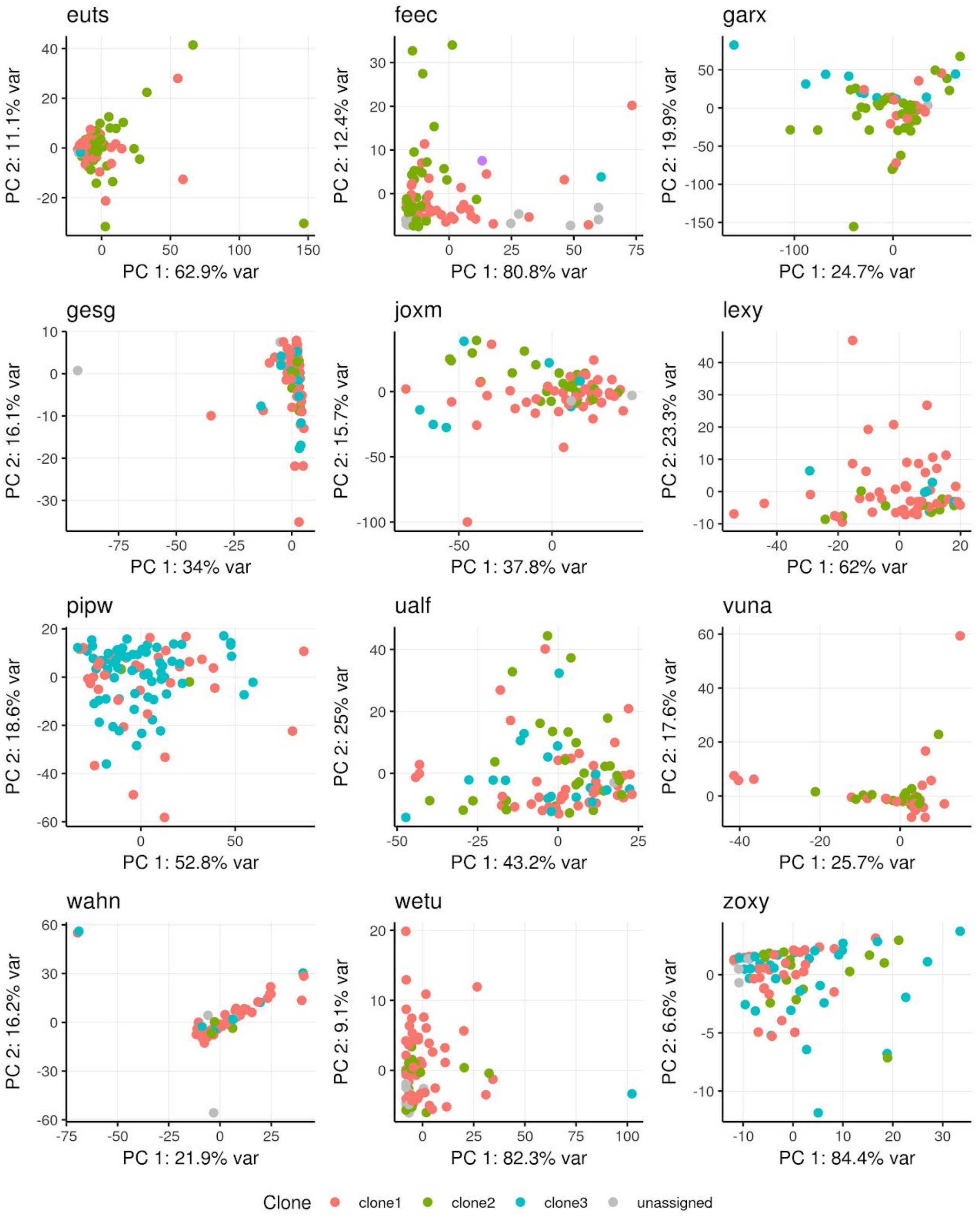


**Fig S9:** Scatter plot of the first two principal components calculated on the read coverage of the set of somatic variant sites used for clone assignment. Shown are data from twelve lines with at least 60 assignable cells. The first two PCs do not segregate cells from different clones, suggesting that read coverage of somatic variants does not associate with or bias clone assignment.

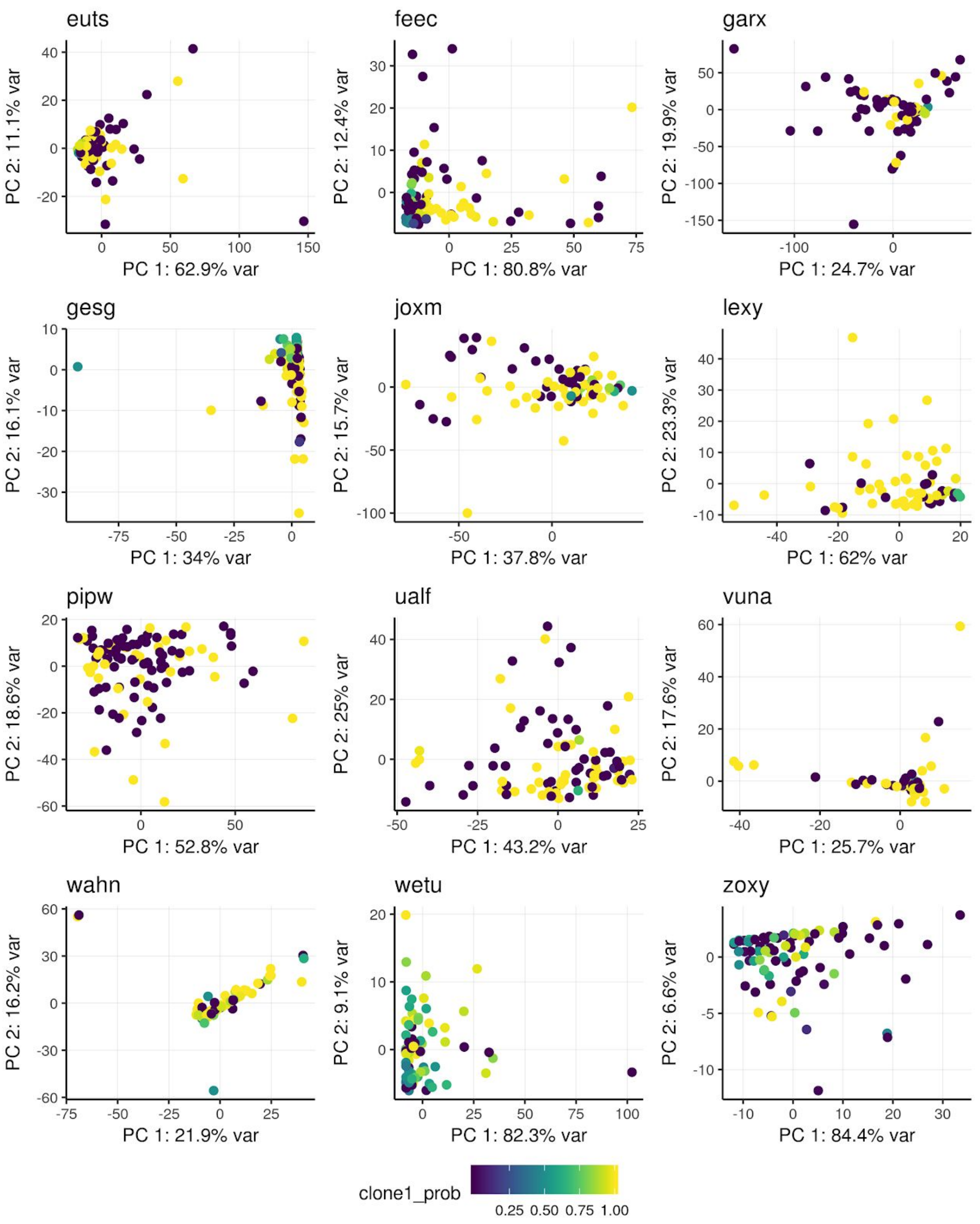


**Fig S10:** Scatter plot of the first two principal components calculated on the read coverage of the set of somatic variant sites used for clone assignment. Cells are colored by the assignment probability of clone 1 (*i.e.* the “base clone” which by definition contains no unique somatic variants). Shown are data from twelve lines with at least 60 assignable cells.

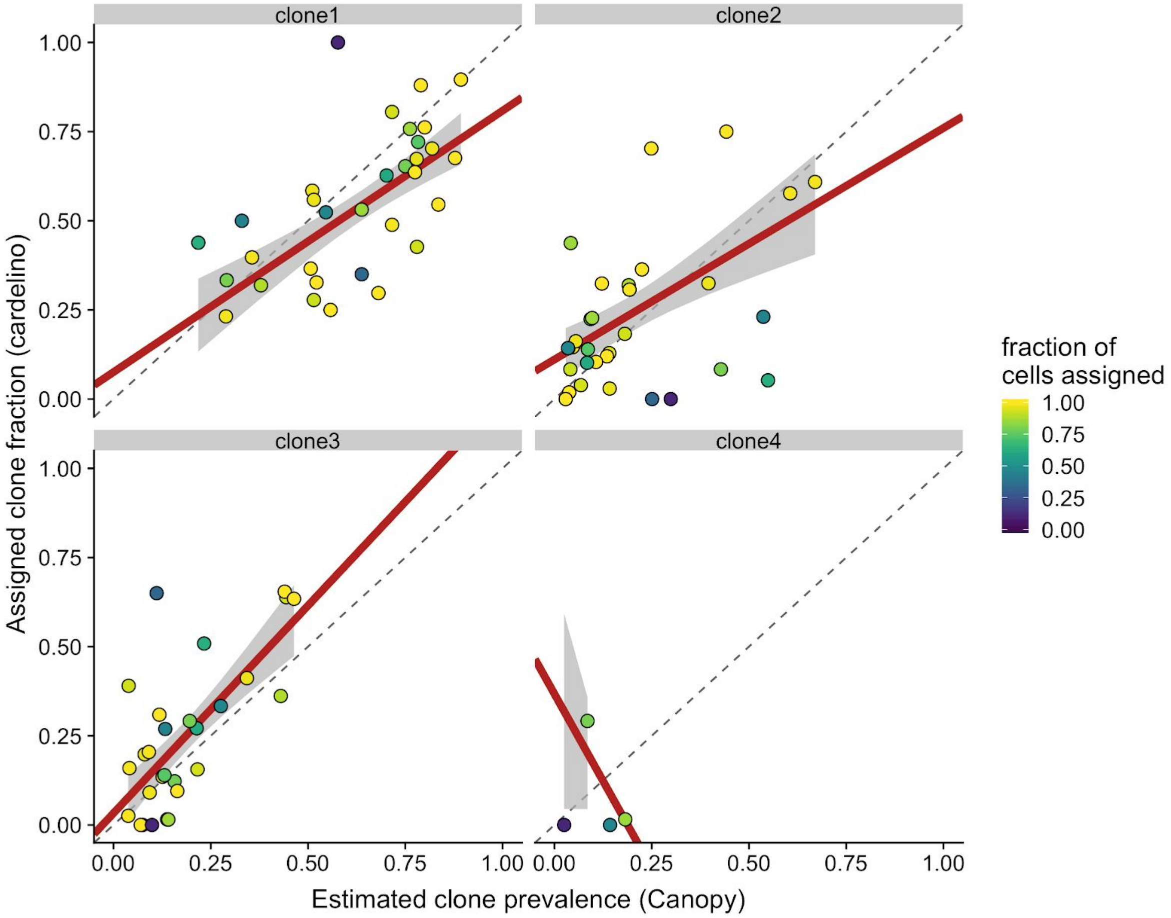


**Fig. S11:** Clone prevalence estimates from WES data (x-axis; using Canopy) *versus* the fraction of single-cell transcriptomes assigned to the clone (y-axis; using *cardelino*), for each clone across lines. Points are coloured by the overall fraction of single-cell transcriptomes assigned for a given line (*i.e.* cells with posterior P>0.5 for assignment).

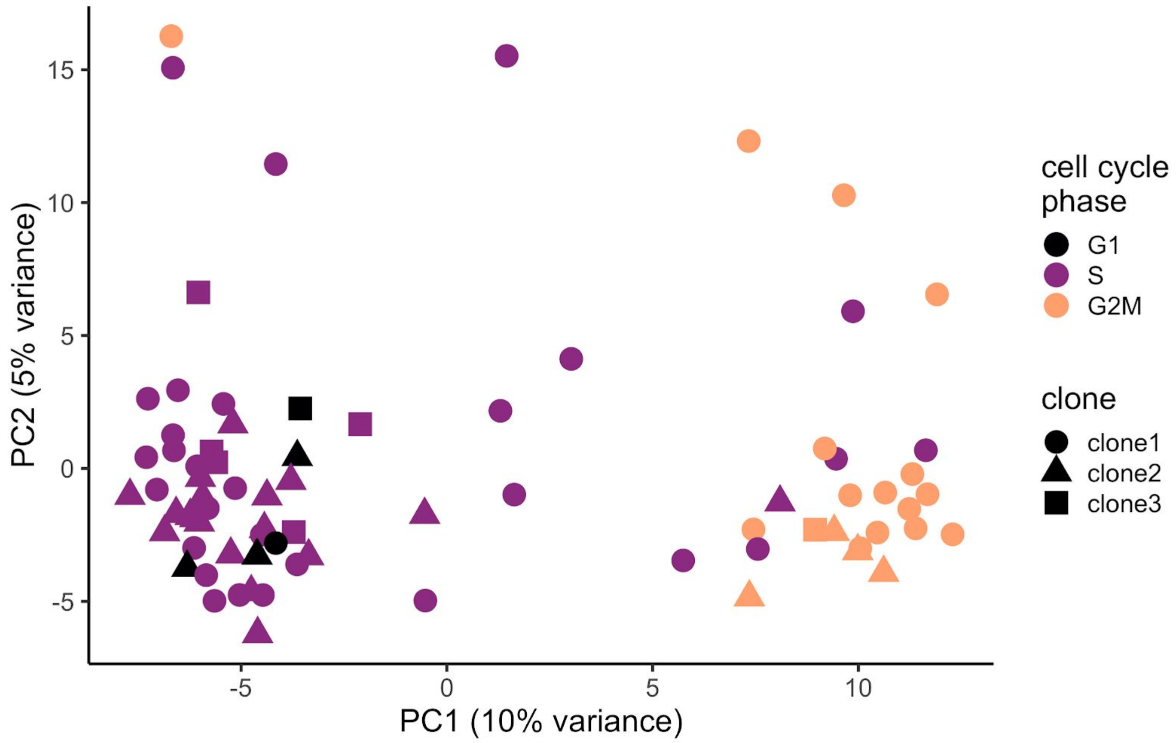


**Fig S12:** Principal component analysis from single-cell gene expression data (top 500 most-variable genes) for clone-assigned cells for the example line *joxm*. Cells are coloured by the cell cycle phase inferred by the cyclone method implemented in the scran package, and shape denotes the assigned clone from cardelino.

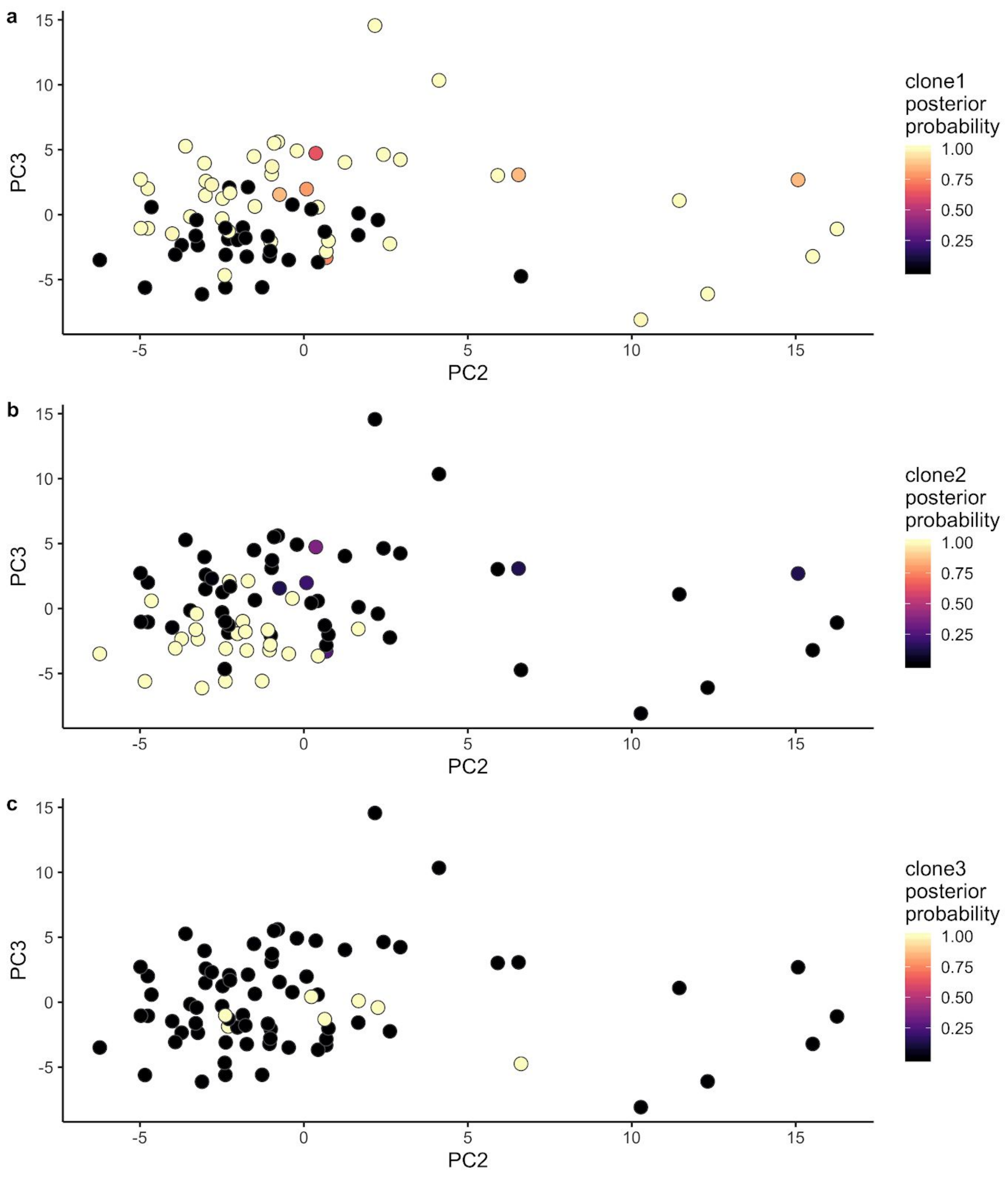


**Fig S13:** Principal component analysis from single-cell gene expression data (top 500 most-variable genes) for clone-assigned cells for the donor *joxm*, plotting principal component 3 against principal component 2. Cells are coloured by the posterior probability from cardelino that the cell belongs to clone1 **(a)**, clone2 **(b)** or clone3 **(c)**.

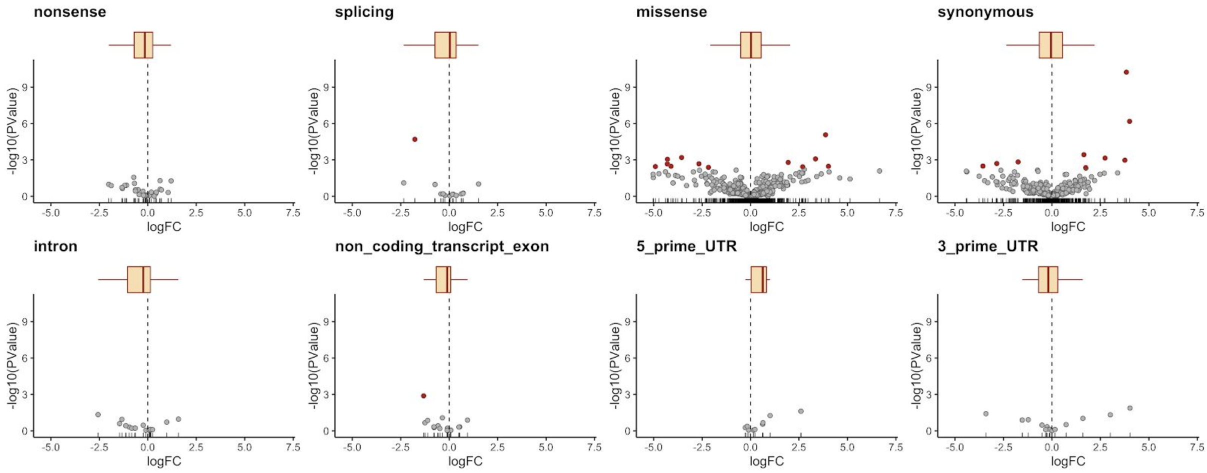


**Fig S14:** Direct effects of somatic variants on genes overlapping the variant. Volcano plot showing -log10(P-value) against $\log_2$-fold change from testing differential expression for genes with a somatic mutation between cells with the mutation and cells without the mutation, faceted by VEP annotation category (**Methods**). Each point represents a gene, and boxplots show the overall $\log_2$-fold change distribution for each annotation category. DE tests are conducted within each line (donor) separately, and results shown here are aggregated across 32 lines. Genes are categorised by simplified functional annotations from VEP of the somatic mutation, and genes significantly DE at an FDR threshold of 20% are shown in red.

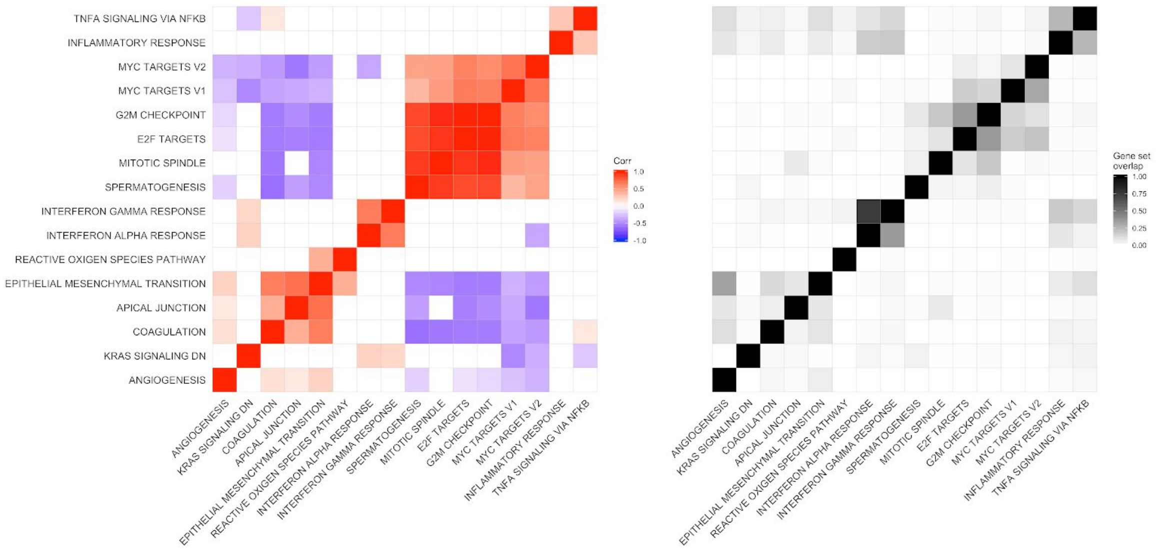


**Fig S15: (left)** Heatmap showing Spearman correlation between gene set enrichment results for the 16 most frequently enriched MSigDB Hallmark gene sets across 31 lines. Colour indicates the correlation between pairs of gene sets and is only shown if the correlation is significant ($P < 0.05$). (**right**) Heatmap showing proportion of overlap in genes between pairs of gene sets (matching those in left panel).

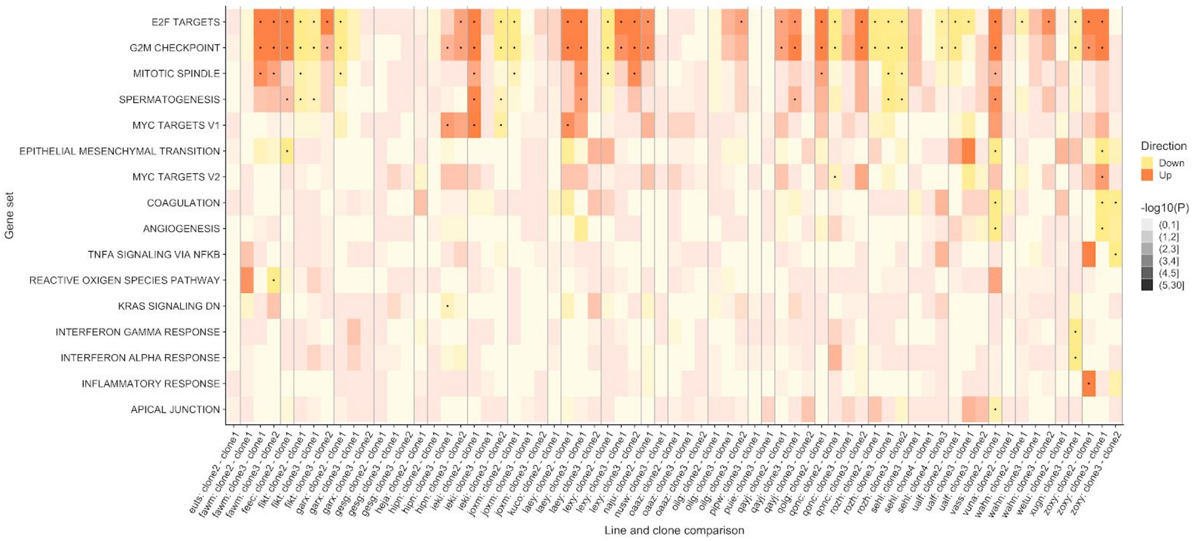


**Fig. S16:** Heatmap showing the direction (first listed clone relative to second listed clone; in colour) and strength of enrichment (-log10(P) as degree of shading) for Hallmark gene sets tested with camera (Methods) for all pairwise comparisons between clones across 31 lines. Gene sets that are significantly enriched at an FDR threshold of 5% are indicated with dots. Gene sets are shown if significant in at least one line, and are ordered by number of lines in which they are significant.

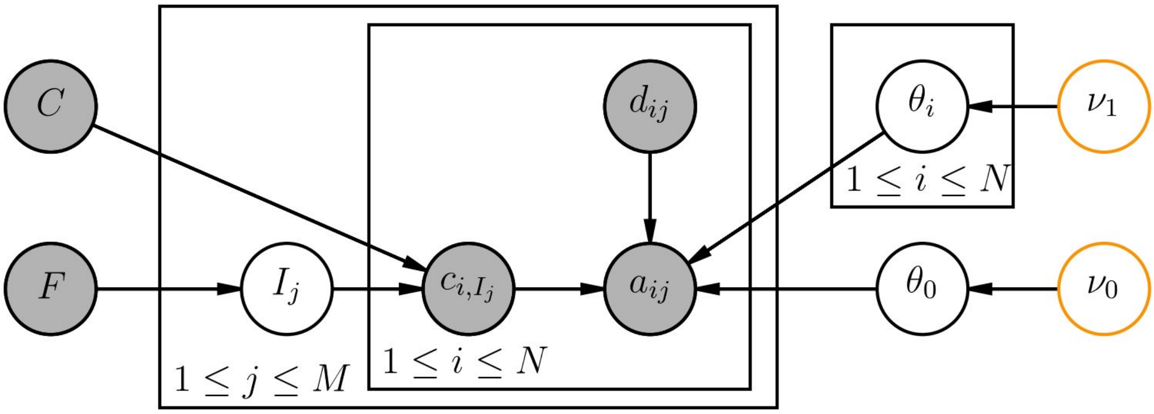


**Fig S17.** Graphical representation of the *cardelino* model. The clonal tree configuration matrix $C$ and clonal fraction *F* are inferred from bulk or single-cell DNA-seq data using existing methods (*e.g. Canopy*) and treated as data by the model. The cell clonal identity $I$ is an unknown variable vector, which together with $C$ encodes the genotype $c_{i,Ij}$ of each variant $i$ in each cell $j$. If $c_{i,Ij}$ is 1, the alternative allelic read count will follow a binomial distribution with gene specific parameter $\theta_i$, otherwise with error related parameter $\theta_0$. Both $\theta_i$ and $\theta_0$ have a beta prior distribution, but with different parameters. Shaded nodes represent observed variables; unshaded nodes represent unknown variables.

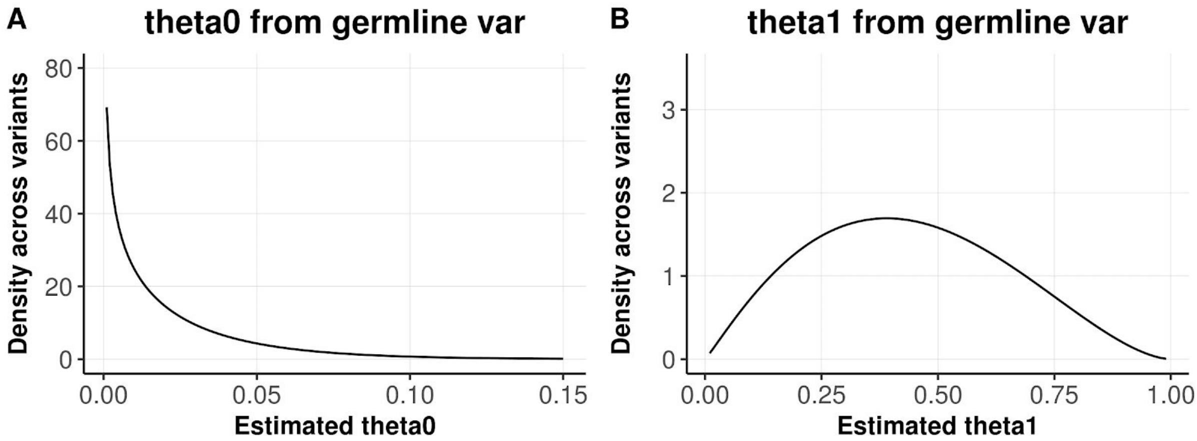


**Fig S18.** Estimated distribution of the "sequencing error rate" (theta0; **A**) and the binomial rate parameter for the alternative allele count given a variant is present (theta1; **B**) from 591 germline heterozygous variants. Each variant has total expressed read counts from 2,000 to 100,000 across 428 cells, and on average has 19.6 read counts per cell. Theta0 is estimated by comparing other bases from genotyped ALT and REF bases, giving a beta distribution of (0.3, 29.7). Theta1 is estimated by comparing the ALT to REF bases, and giving beta distribution of (2.25, 2.65). Format of beta distribution parameters: (shape1, shape2).

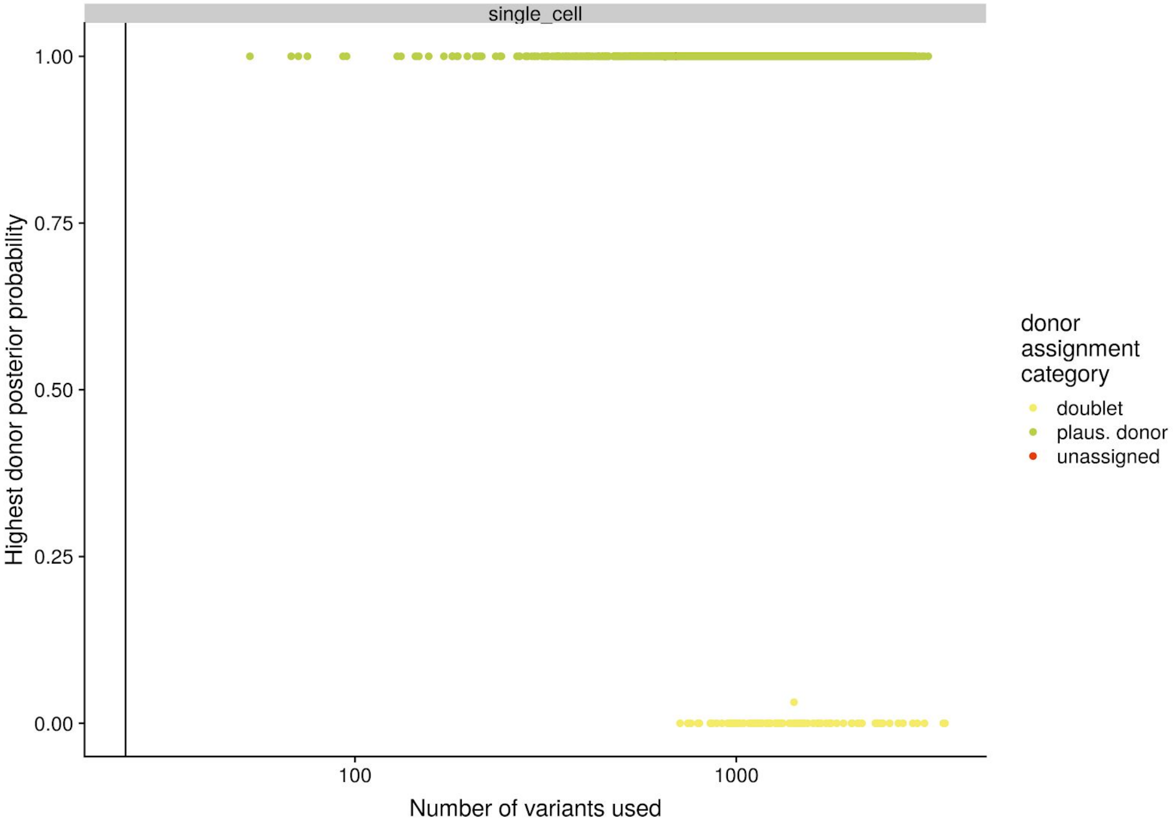


**Fig S19:** Donor identification results from cardelino for QC-passing cells for 32 fibroblast lines (*i.e.* donors) used to demultiplex cells from plates on which cells from three lines were pooled. The y-axis shows the highest posterior probability for donor assignment from cardelino (**Methods**) for a little over 2,000 cells passing QC using expression-based metrics (real Smart-seq2 data from our study; not simulated data). The donor ID results are emphatic, with posterior probabilities either very close to 1 or very close to zero, meaning that the model is very confident about assigning each cell either to a specific donor (*i.e.* line) or that the “cell” is actually doublet, or that it matches none of the plausible donors. The x-axis shows the number of germline variants with read coverage in the cells that were informative for donor assignment of the cell. Cells are coloured by donor assignment category: either “plausible donor” *(i.e.* a donor/line that was known to have been used on the processing plate), “doublet” (nominal single cells that have been inferred to be doublets) or “unassigned” (too few variants for assignment or posterior probability of assignment less than 0.95). NB: 21 unassigned cells are not visible due to overplotting by doublet cells.

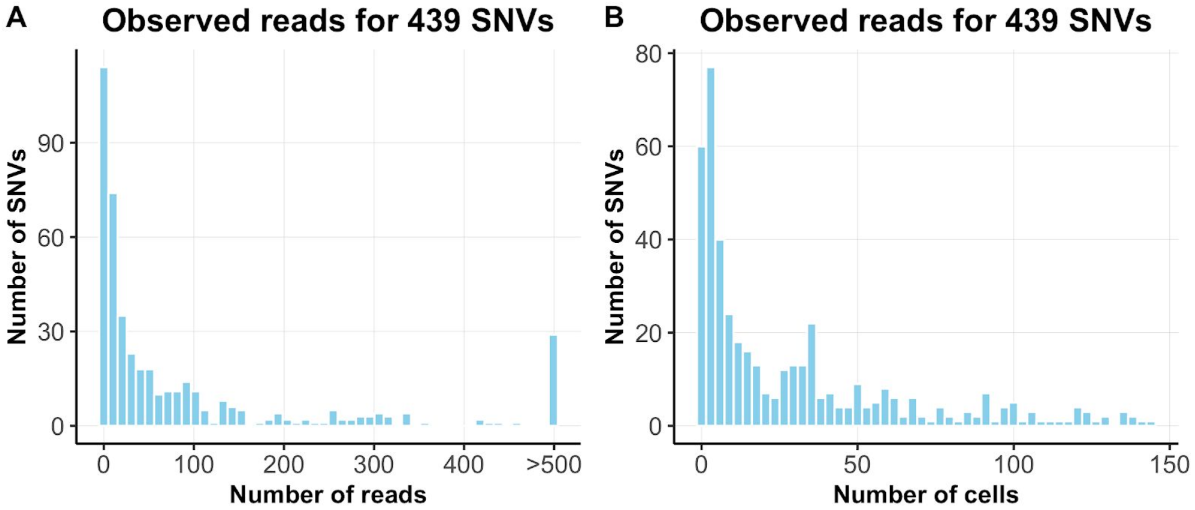


**Fig S20.** Summary of sequencing depths of 439 variants (single nucleotide variants, SNVs) across a pool of 151 cells. **(A)** Histogram of total read counts on each variant; **(B)** Histogram of the number of cells with non-zero read coverage for each variant. This matrix is used as a seed to generate sequencing depths for simulations in Fig 1(b-d) and Fig S1.
