## Supplementary material for "Cardelino: Integrating whole exomes and single-cell transcriptomes to reveal phenotypic impact of somatic variants"

#### 1 The cardelino model

As input for cardelino, we assume that a clonal structure configuration is first inferred from deep exome-seq, for example using Canopy [1]. This yields clonal fractions $F = (f_1, .., f_K)$, where $f_k$ denotes the relative prevalence of a given clone $k$ ($\sum_{k=1}^{K} f_k = 1$), as well as a clonal tree configuration matrix $C$ (an $N$-by-$K$ binary matrix) for $N$ variants and $K$ clones, where $c_{i,k} = 1$ if somatic variant $i$ is present in clone $k$ and $c_{i,k} = 0$ otherwise. Given $C$ and $F$, cardelino is aimed at assigning individual cells to one of $K$ clones based on their expressed alleles using a probabilistic clustering model (see graphical representation in **Supp. Fig. S21**). Based on scRNA-seq, we extract for each cell and variant that segregates between clones the number of sequencing reads supporting the reference allele (reference read count) and the number of reads supporting the alternative allele (alternate read count). We denote the variant-by-cell matrix of alternate read counts by $A$ and the variant-by-cell matrix of total read counts (sum of reference and alternate read counts) by $D$. Entries in $A$ and $D$ are therefore non-negative integers, with missing entries in the matrix a indicating zero read coverage for a given cell and variant.

The prior probability that cell $j$ belongs to clone $k$ could be taken as the clonal fraction $f_k$, but to avoid biasing cell assignment towards highly prevalent clones for cells with little read information (where the prior is more influential) we use a uniform prior $F$ such that $P(I_j = k|F) = 1/K$ for all $k$. Note, the variable $F$ is used to denote a uniform prior for convenience here, which can be different from Canopy output. Given this prior distribution, the posterior probability of cell $j$ belonging to clone $k$ can be expressed as:

$$P(I_j = k|\boldsymbol{a_j}, \boldsymbol{d_j}, C, F, \boldsymbol{\theta}) = \frac{P(\boldsymbol{a_j}|\boldsymbol{d_j}, I_j = k, C, \boldsymbol{\theta})P(I_j = k|F)}{\sum_{t=1}^{K} P(\boldsymbol{a_j}|\boldsymbol{d_j}, I_j = t, C, \boldsymbol{\theta})P(I_j = t|F)}, \tag{1}$$

where $I_j$ is the identity of the specific clone cell $j$ is assigned to, and $\boldsymbol{a_j}$ and $\boldsymbol{d_j}$ are the observed alternate read count and total read count vectors, respectively, for variants 1 to $N$ in cell $j$. The parameter vector $\boldsymbol{\theta}$ is a set of unknown parameters to model the allelic counts, which will be discussed in next section.

#### 2 Modelling allelic imbalance

The core part of the cardelino model is to model the alternate read count using a binomial model. For a given site in a given cell, there are two possibilities: the variant is "absent" in the clone a cell is assigned to (i.e. the cell is homozygous reference at that position) or the variant is “present” in the clone the cell is assigned to (i.e. the cell is heterozygous at that position), as encoded in the configuration matrix $C$. When considering the “success probability” $\boldsymbol{\theta}$ for the binomial model, where here a success is defined as observing an alternate read, we consider two alternative (sets of) parameters for each of these settings: $\theta_0$ for homozygous reference alleles (variant absent), and $\boldsymbol{\theta_1} = \{\theta_1, ..., \theta_N\}$ for the case with heterozygous variants (variant present). Note, here we use a common parameter $\theta_0$ for homozygous reference alleles in all variants, but $\theta_i, i \geq 1$ for each variant $i$ to account for the gene specific level of allelic imbalance that causes the probability of observing alternate reads to differ from 0.5. Therefore, the allelic counts base model for the two genotypes can be written in the following binomial distributions,

$$p(a_{i,j}|d_{i,j}, h_{i,j}, \boldsymbol{\theta}) = \begin{cases} \texttt{Binom}(a_{i,j}|d_{i,j}, \theta_0), & \text{if } h_{i,j} = 0. \\ \texttt{Binom}(a_{i,j}|d_{i,j}, \theta_i), & \text{if } h_{i,j} = 1. \end{cases} \tag{2}$$

where $h_{i,j} = c_{i,I_j} \in \{0, 1\}$ is the genotype of variant $i$ in cell $j$, which is encoded by clonal configuration $C$ and cell identity $I_j$. Furthermore, the likelihood of cell $j$ from clone $k$ can be formalised as follows,

$$\begin{aligned} P(\boldsymbol{a_j}|\boldsymbol{d_j}, I_j = k, C, \boldsymbol{\theta}) &= \prod_{i=1}^{N} p(a_{i,j}|d_{i,j}, h_{i,j}, \boldsymbol{\theta}) \\ &= \prod_{i=1}^{N} \left\{ \texttt{Binom}(a_{i,j}|d_{i,j}, \theta_i)^{c_{i,k}} \times \texttt{Binom}(a_{i,j}|d_{i,j}, \theta_0)^{1-c_{i,k}} \right\} \end{aligned} \tag{3}$$

Then, we could have the likelihood of parameters $\boldsymbol{\theta} = \{\theta_0, \theta_1, ..., \theta_N\}$ to observe a full data set across $M$ cells by marginalizing the mixture of cell assignment, as follows

$$\mathcal{L}(\boldsymbol{\theta}) = \prod_{j=1}^{M} \sum_{I_j=1}^{K} P(\boldsymbol{a_j}|\boldsymbol{d_j}, I_j, C, \boldsymbol{\theta}) P(I_j|F). \tag{4}$$

Furthermore, we could view the the clonal assignment in a Bayesian way, and introduce informative prior distributions for unknown parameters $\boldsymbol{\theta}$. By multiplying the prior probability by the likelihood, we could have the posterior probability as follows,

$$\begin{aligned} P(\boldsymbol{\theta}|A, D, C, F, \boldsymbol{\nu}) &\propto P(\boldsymbol{\theta}|\boldsymbol{\nu}) \times \prod_{j=1}^{M} \sum_{I_j=1}^{K} P(\boldsymbol{a_j}|\boldsymbol{d_j}, I_j, C, \boldsymbol{\theta}) P(I_j|F) \\ &= \texttt{Beta}(\theta_0|\alpha_0, \beta_0) \prod_{i=1}^{N} \texttt{Beta}(\theta_i|\alpha_1, \beta_1) \times \prod_{j=1}^{M} \sum_{I_j=1}^{K} P(\boldsymbol{a_j}|\boldsymbol{d_j}, I_j, C, \boldsymbol{\theta}) P(I_j|F), \end{aligned} \tag{5}$$

where we use a beta prior distribution, a conjugate distribution to binomial distribution, for each $\theta$, and the hyperparameters $\boldsymbol{\nu} = \{\alpha_0, \beta_0, \alpha_1, \beta_1\}$ of prior are learned from germline heterozygous variants.

Accounting for the uncertainty of $\boldsymbol{\theta}$, this unknown parameter can be marginalised in the posterior probability of clonal assignment, as follows,

$$P(I_j = k|\boldsymbol{a_j}, \boldsymbol{d_j}, C, F) = \int_{\boldsymbol{\theta}} P(I_j = k|\boldsymbol{a_j}, \boldsymbol{d_j}, C, F, \boldsymbol{\theta}) P(\boldsymbol{\theta}|A, D, C, F, \boldsymbol{\nu}) d\boldsymbol{\theta}. \tag{6}$$

### 3 Inference for the cardelino model

In the above section, we defined the posterior probability of clonal assignment and unknown parameter $\boldsymbol{\theta}$. With conjugate prior distributions, a Gibbs sampler can be used to generate a set of samples following the posterior distribution.

In this Gibbs sampling algorithm, we sample cell assignment $I$ and parameters $\boldsymbol{\theta}$ alternately. Given that one of these two unknown variables is fixed, the elements of the other parameter are conditionally independent. Therefore, given the parameters $\boldsymbol{\theta}$, we could sample the clonal identity $I_j$ via a categorical distribution, taking Eq(3,2), as follows

$$\begin{aligned} P(I_j = k | I_{-j}, A, D, C, F, \boldsymbol{\theta}) &= P(I_j = k | \boldsymbol{a_j}, \boldsymbol{d_j}, C, F, \boldsymbol{\theta}) \\ &\propto P(I_j = k | F) P(\boldsymbol{a_j} | I_j = k, \boldsymbol{d_j}, C, \boldsymbol{\theta}). \end{aligned} \tag{7}$$

Similarly, given the clonal identity $I$ in a previous step, $\theta_i, 0 \leq i \leq N$ are independent from each other, and the posterior probability in Eq(5) can be rewritten by inserting the base model in Eq(2) as follows,

$$\begin{aligned} P(\boldsymbol{\theta} | A, D, C, I, \boldsymbol{\nu}) &\propto \texttt{Beta}(\theta_0 | \alpha_0, \beta_0) \prod_{i=1}^{N} \texttt{Beta}(\theta_i | \alpha_1, \beta_1) \\ &\times \prod_{j=1}^{M} \prod_{i=1}^{N} \texttt{Binom}(a_{i,j} | d_{i,j}, \theta_0)^{1 - c_{i,I_j}} \texttt{Binom}(a_{i,j} | d_{i,j}, \theta_i)^{c_{i,I_j}} \\ &= \texttt{Beta}(\theta_0 | \alpha_0, \beta_0) \prod_{j=1}^{M} \prod_{i=1}^{N} \texttt{Binom}(a_{i,j} | d_{i,j}, \theta_0)^{1 - c_{i,I_j}} \\ &\times \prod_{i=1}^{N} \left\{ \texttt{Beta}(\theta_i | \alpha_1, \beta_1) \prod_{j=1}^{M} \texttt{Binom}(a_{i,j} | d_{i,j}, \theta_i)^{c_{i,I_j}} \right\}. \end{aligned} \tag{8}$$

Therefore, for individual $\theta$, we could sample it via a beta distribution as follows,

$$\theta_0 | I \sim \texttt{beta}(\alpha_0 + u_0, \beta_0 + v_0); \qquad \theta_i | I \sim \texttt{beta}(\alpha_1 + u_i, \beta_1 + v_i), i > 0 \tag{9}$$

where

$$\begin{aligned} u_0 &= \sum_{i=1}^{N} \sum_{j=1}^{M} a_{i,j}(1 - c_{i,I_j}), \qquad & v_0 &= \sum_{i=1}^{N} \sum_{j=1}^{M} (d_{i,j} - a_{i,j})(1 - c_{i,I_j}), \\ u_i &= \sum_{j=1}^{M} a_{i,j} c_{i,I_j}, \;\; i > 0, \qquad & v_i &= \sum_{j=1}^{M} (d_{i,j} - a_{i,j}) c_{i,I_j}, \;\; i > 0. \end{aligned} \tag{10}$$

Now, based on Eq (7, 9), we could sample the full joint distribution of $I$ and $\theta$ with Gibbs sampling in the following Algorithm 1.

**Algorithm 1:** Gibbs sampling for cell assignments to clones

```
Initialize θ = {θ_0, θ_1, ..., θ_N}
for t = 1 to H do
    for j = 1 to M do
        Sample: I_j = k | I_{-j}, A, D, C, F, θ with Eq(7)
    for i = 0 to N do
        Sample: θ_i | I, A, D, C, θ_{-i} with Eq (9)
```

In practice, we could sample 1,000 iterations and check the convergence with Geweke's convergence diagnostic ($Z$ score) by using the first 10% and the last 50% iterations of the sampled chain. If $|Z| > 2$, then 100 more iterations will be added until the criterion is passed. Usually, this algorithm converges very quickly, even with as few as 100 iterations in some cases.

### 4 Alternative models and estimation with EM algorithm

Besides the cardelino model, we also compared with two alternative models, where we assume all sites have a common parameter when variant is "present", i.e., $\theta_1 = \theta_2 = ... = \theta_N$. For simplicity, we use $\theta_1$ to denote this shared parameter and ignore the conflict with the symbol in the cardelino model. Therefore, the alternative models only have two parameters $\theta_0$ and $\theta_1$, for the "success probability" for variant absent and present, respectively. In the simplest model **theta_fixed**, we fix $\theta_0 = 0.01$ and $\theta_1 = 0.5$.

Besides the fixed values, these parameters may be different from experiments to experiments and also vary according to the accuracy of clonal configuration that is inferred in a previous step. Therefore, it is important to fit the parameters to the observed data. A point estimate of these parameter $\boldsymbol{\theta} = \{\theta_0, \theta_1\}$ with maximum likelihood can be achieved with an Expectation-Maximization (EM) algorithm (termed as **theta_EM**).

EM algorithm has the advantage of being much more computationally efficient than the Gibbs sampler. However, the point estimate will lose the uncertainty in the parameters for clonal assignment, and consequently can suffer from over-fitting if there are very few sequencing reads, especially in lowly expressed genes. Therefore, it is important to use a single parameter for all variants and turn off the gene specific parameters in original Eq(2) to retain sufficient reads for a robust point estimate. This setting can be very useful in assigning cells to donors given genotypes in multiplexed experiments, where the statistical framework is the same but the error in the genotypes is much lower than from a clonal tree, and the large number of variants could benefit from the high efficiency of the EM algorithm. Here, we introduce the algorithm with all $\theta_i, 1 \leq i \leq N$ turned into a single shared parameter $\theta_1$; all equations in above sections still hold.

In order to maximise the likelihood in Eq(4) (or log likelihood for convenience), let us first rewrite the likelihood of assigning a single cell $j$ to a certain clone $k$ by extending the binomial probability as follows,

$$\begin{aligned} P(\boldsymbol{a_j}|\boldsymbol{d_j}, I_j = k, C, \boldsymbol{\theta}) &= \prod_{i=1}^{N} P(a_{i,j}|d_{i,j}, \theta, c_{i,k}) = \prod_{i=1}^{N} \mathcal{B}(a_{i,j}; d_{i,j}, \theta_{c_{i,k}}) \\ &= w_j \times \theta_0^{S^1_{j,k}} \times (1-\theta_0)^{S^2_{j,k}} \times \theta_1^{S^3_{j,k}} \times (1-\theta_1)^{S^4_{j,k}}, \end{aligned} \tag{11}$$

where $w_j = \prod_{i=1}^{N} \binom{d_{i,j}}{a_{i,j}}$ is a product of binomial coefficients. $S^1_{j,k}, S^2_{j,k}, S^3_{j,k}, S^4_{j,k}$ are the summarized read counts of alternative and reference alleles in genotypes without or with variant, respectively, as follows,

$$\begin{aligned} S^1_{j,k} &= \sum_{i=1}^{N} a_{i,j}\mathbb{I}(c_{i,k}=0), \qquad S^2_{j,k} = \sum_{i=1}^{N} (d_{i,j} - a_{i,j})\mathbb{I}(c_{i,k}=0), \\ S^3_{j,k} &= \sum_{i=1}^{N} a_{i,j}\mathbb{I}(c_{i,k}=1), \qquad S^4_{j,k} = \sum_{i=1}^{N} (d_{i,j} - a_{i,j})\mathbb{I}(c_{i,k}=1). \end{aligned} \tag{12}$$

These values can be equivalently taken from dot products of matrices $S^1 = A^\top(1-C)$, $S^2 = (D-A)^\top(1-C)$, $S^3 = A^\top C$, and $S^4 = (D-A)^\top C$.

Now, we can estimate the clonal assignment $I_j$ and the parameters $\boldsymbol{\theta} = \{\theta_0, \theta_1\}$ with an EM algorithm. In the initialization, we set the parameter $\theta$ randomly. Then we iterate the E step and M step in the EM algorithm. In the E-step, given the parameter in the previous step, we calculate the posterior of the cell assignment

$$\gamma_{j,k} = P(I_j = k|\boldsymbol{a_j}, \boldsymbol{d_j}, C, F, \theta) = \frac{P(\boldsymbol{a_j}|\boldsymbol{d_j}, I_j = k, C, \boldsymbol{\theta})P(I_j = k|F)}{\sum_{t=1}^{K} P(\boldsymbol{a_j}|\boldsymbol{d_j}, I_j = t, C, \boldsymbol{\theta})P(I_j = t|F)}, \tag{13}$$

which is often called component responsibility in the EM algorithm. In the M-step, given the posterior of cell assignment, we optimize the parameter to maximize the likelihood. By setting the derivation of the log likelihood Eq (4) (taking Eq (11)) to 0, we could have the following condition to satisfy,

$$\frac{\log \mathcal{L}(\theta)}{\theta_0} = \sum_{j=1}^{M}\sum_{k=1}^{K} \gamma_{j,k} \left[ \frac{S_{j,k}^1}{\theta_0} - \frac{S_{j,k}^2}{1-\theta_0} \right] = 0. \tag{14}$$

Therefore, we can have a closed form solution for $\theta_0$ (and $\theta_1$ similarly) as follows,

$$\theta_0 = \frac{\sum_{j=1}^{M}\sum_{k=1}^{K} \gamma_{j,k} S_{j,k}^1}{\sum_{j=1}^{M}\sum_{k=1}^{K} \gamma_{j,k} (S_{j,k}^1 + S_{j,k}^2)} \qquad \theta_1 = \frac{\sum_{j=1}^{M}\sum_{k=1}^{K} \gamma_{j,k} S_{j,k}^3}{\sum_{j=1}^{M}\sum_{k=3}^{K} \gamma_{j,k} (S_{j,k}^3 + S_{j,k}^4)}. \tag{15}$$

Here, we summarize the EM algorithm for the cell assignment and parameter estimate in the following Algorithm 2. To end the algorithm, we could check if the improvement of the log likelihood is lower than a threshold or set a fixed number of iterations (e.g. 100 iterations are sufficient in many cases).

---

**Algorithm 2:** EM algorithm for cell assignments to clones

---

**Initialize** $\theta = \{\theta_0, \theta_1\}$ and evaluate $\log \mathcal{L}(\theta)$

**while** *not converged* **do**

    **E step**: Calculate $\gamma_{j,k}$ with current parameters

    $\gamma_{j,k} = \frac{P(A_j|I_j=k,D_j,C,F,\theta)P(I_j=k)}{\sum_{t=1}^{K} P(A_j|I_j=t,D_j,C,F,\theta)P(I_j=t)}$

    **M step**: Maximizing likelihood on parameters with current responsibilities

    $\theta_0^{\texttt{new}} = \frac{\sum_{j=1}^{M}\sum_{k=1}^{K} \gamma_{j,k} S_{j,k}^1}{\sum_{j=1}^{M}\sum_{k=1}^{K} \gamma_{j,k} (S_{j,k}^1 + S_{j,k}^2)}; \quad \theta_1^{\texttt{new}} = \frac{\sum_{j=1}^{M}\sum_{k=1}^{K} \gamma_{j,k} S_{j,k}^3}{\sum_{j=1}^{M}\sum_{k=3}^{K} \gamma_{j,k} (S_{j,k}^3 + S_{j,k}^4)}$

    **Update** $\log \mathcal{L}(\theta)$ and check convergence

**return** $\theta, \boldsymbol{\gamma}, \log \mathcal{L}(\theta)$

---

In addition, the binomial distribution can be switched into simpler Bernoulli model by setting a threshold $s$ (e.g. 1) as $\hat{a}_{i,j} = \mathbb{I}(a_{i,j} \geq s)$ and $\hat{d}_{i,j} = \mathbb{I}(d_{i,j} \geq s)$, and all above equations and inference methods remain applicable. The Bernoulli base model can be useful when the sequencing coverage is highly even, e.g., in scDNA-seq [2] or when the variance of allelic expression is extremely high.
